# Systemic identification of functionally conserved lncRNA metabolic regulators in human and mouse livers

**DOI:** 10.1101/2024.08.10.607444

**Authors:** Chengfei Jiang, Zhe Li, Ping Li, Yonghe Ma, Sunmi Seok, Stephanie K. Podguski, Shria Moturi, Nao Yoneda, Kenji Kawai, Shotaro Uehara, Yasuyuki Ohnishi, Hiroshi Suemizu, Jinwei Zhang, Haiming Cao

## Abstract

**BACKGROUND & AIMS:** Unlike protein-coding genes, the majority of human long non-coding RNAs (lncRNAs) lack conservation based on their sequences, posing a challenge for investigating their role in a pathophysiological context for clinical translation. This study explores the hypothesis that non-conserved lncRNAs in human and mouse livers may share similar metabolic functions, giving rise to functionally conserved lncRNA metabolic regulators (fcLMRs).

**METHODS:** We developed a sequence-independent strategy to select putative fcLMRs, and performed extensive analysis to determine the functional similarities of putative human and mouse LMR pairs (h/mLMRs).

**RESULTS:** We found that several pairs of putative fcLMRs share similar functions in regulating gene expression. We further demonstrated that a pair of fcLMRs, h/mLMR1, robustly regulated triglyceride levels by modulating the expression of a similar set of lipogenic genes. Mechanistically, h/mLMR1 binds to PABPC1, a regulator of protein translation, via short motifs on either lncRNA with divergent sequences but similar structures. This interaction inhibits protein translation, activating an amino acid-mTOR-SREBP1 axis to regulate lipogenic gene expression. Intriguingly, PABPC1-binding motifs on each lncRNA fully rescued the functions of their corresponding LMRs in the opposite species. Given the elevated expression of h/mLMR1 in humans and mice with hepatic steatosis, the PABPC1-binding motif on hLMR1 emerges as a potential non-conserved human drug target whose functions can be fully validated in a physiologically relevant setting before clinical studies.

**CONCLUSIONS:** Our study supports that fcLMRs represent a novel and prevalent biological phenomenon, and deep phenotyping of genetic mLMR mouse models constitutes a powerful approach to understand the pathophysiological role of lncRNAs in the human liver.

## Introduction

Elucidating the intricate molecular circuitrie governing metabolic regulation in the human liver is imperative for a comprehensive understanding of the pathophysiological underpinnings of metabolic diseases and the development of effective therapeutic interventions. Although the regulatory networks of liver metabolism are well defined at the protein-coding gene level, there remains a significant gap in understanding the role of long non-coding RNAs (lncRNAs) in the human liver. LncRNAs constitute the largest transcript class of the human genome^1, 2^ and the functions of lncRNAs has been integrated into all major cell processes^3, 4^. Although genetic and clinical data have connected a growing number of human lncRNAs to the pathogenesis of metabolic disorders^5–7^, the metabolic functions of most human lncRNAs in the liver are largely unknown, owing to the inability to define their role in a physiological context. Critical regulators of energy metabolism often exert their function by coordinating the actions of multiple organs; defining their role in metabolism always requires careful examination of their function in a physiological context. For protein coding genes, this has been routinely carried out in animal models such as mice. This approach, however, is uniquely difficult for human lncRNA genes. Unlike protein coding genes, most human lncRNAs are not conserved, even among other mammals^3, 8^, restricting the ability to study their physiological function in animal models. Therefore, despite growing evidence supporting the role of human lncRNAs in the pathogenesis of human metabolic diseases, well-characterized human lncRNA metabolic regulators (hLMRs) are still scarce; and the inability to vigorously define their in vivo role is likely to be one of the leading confounding factors.

Although most human lncRNAs are considered non-conserved if analyzed by current sequence comparison tools such as BLAST, additional possibilities regarding their conservation indeed exist as our current understanding of the sequence-function relationship of lncRNAs remains extremely limited. For example, if some lncRNAs are conserved at the level of small functional RNA motifs or 3D structure, they will never be identified by current sequence-based tools. If such lncRNAs indeed exist, defining functionally conserved lncRNA metabolic regulators (fcLMRs) in the human liver could fundamentally change the way that we study the pathophysiological role of lncRNAs and rapidly increase the overall understanding of human liver physiology. But as discussed above, it is currently very challenging to define the physiological role of any potential hLMRs, let alone to compare their functions with lncRNAs in other species to identify fcLMRs. To address this critical question at a systematic level, we developed a novel strategy to screen for lncRNA pairs in human and mouse livers that have best potential to function as fcLMRs. More importantly, we performed rigorous validation of our selection by simultaneously confirming multiple pairs of putative fcLMRs to be legitimate ones. Our in-depth analysis of one fcLMR pair also demonstrated that the hLMR and its corresponding mLMR can bind to the same protein through which they engaged essential metabolic pathways, i.e., SREBP1 and mTOR, to regulate metabolic flux. Our results support that fcLMRs are a novel concept of lncRNA biology that can be leveraged to accelerate the delineation of disease-associated hLMRs in the human liver and facilitate their translation to therapy development efforts in humans.

## Results

### A function-based strategy to identify functionally conserved lncRNA metabolic regulators (fcLMRs) in human and mouse livers

To maximize the chance to identify fcLMRs in human and mouse livers, we developed a strategy to select putative fcLMRs that could have similar functions based on the following three criteria: being syntenic, having similar regulatory responses to metabolic stimuli in chimeric human-mouse livers, and being correlated to the same metabolic pathways in isogenic humanized livers (Figure 1*A*).

**Figure 1.**
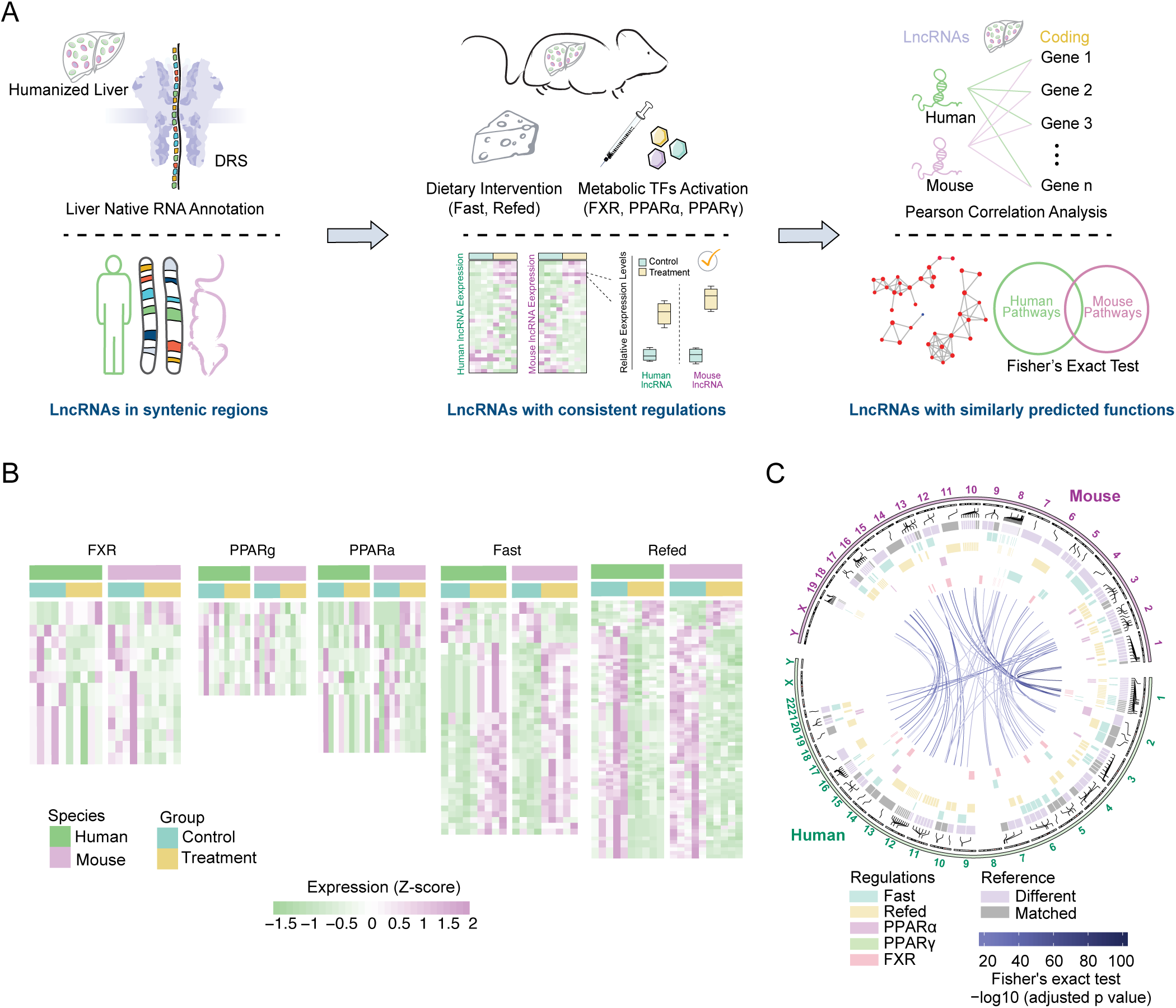
A function-based strategy to identify functionally conserved lncRNA metabolic regulators (fcLMRs) between humans and mice (A) Schematic diagram of the identification of fcLMRs. (B) The heatmaps depict the expression levels of lncRNA pairs that were consistently and significantly regulated (|log2(fold change)|>0.5 and p value<0.05) under the same treatment conditions in both humans and mice. (C) Circular diagram of the genomic locations of potential fcLMRs. Purple markings on the inner circle indicated that the isoform of the lncRNA differed from the reference genome, while gray indicated that they were the same. Information regarding the regulation of the lncRNAs under different treatments was summarized in the center. The LMR pairs were shown in the center with purple lines, and the depth of the line color represented the -log10(adjusted p value).

Evidently, the success of our selection pipeline depends on the accuracy and inclusiveness of lncRNA annotations for human and mouse livers. As lncRNA expression is highly condition-dependent and tissue-specific^3, 9^ and most lncRNA genes produce multiple transcripts, we recently established *de novo* annotations of human and mouse liver lncRNAs at the transcript level under diverse metabolic conditions^10^. Based on the novel annotations, we identified lncRNAs in syntenic regions of human and mouse genomes that are flanked by orthologous protein-coding genes, resulting in 1538 potential fcLMR pairs (Supplementary Figure 1A and Supplementary Table 1). We further filtered the potential fcLMR pairs by their similarity in regulation under physiologically relevant metabolic conditions. When we recently established the novel annotations of human lncRNAs^10^, we employed a chimeric human-mouse liver model in which human and mouse hepatocytes share the same circulation and tissue environment, and subjected the humanized mice to typical metabolic conditions including various feeding conditions (normal chow, 24-hour fast, and 4-hour refeed after 24-hour fast) and transcription factor activations (PPARα, PPARγ, and FXR agonists). Comparing the regulation of human and mouse lncRNAs in the humanized livers led us to identify 118 similarly regulated lncRNA pairs in humans and mice (Figure 1*B* and Supplementary Table 2). Finally, we performed lncRNA-mRNA correlation analyses using gene expression data to infer the lncRNAs’ biological role based on the function of correlated mRNAs. As the diverse genetic backgrounds and environmental exposures of humans could complicate gene-gene interaction networks, we reasoned that RNA-seq data of livers from humanized mice — which are isogenic and housed under the same environment — might perform better in a correlation analysis. Indeed, we found that correlation analysis based on data of humanized mice could more effectively identify orthologous protein coding genes between humans and mice (Supplementary Figure 1B). Therefore, we used data of humanized mice to identify the top protein-coding genes associated with human and mouse lncRNAs. We then conducted pathway enrichment analysis on these correlated coding genes to infer the biological pathways and functions in which the lncRNAs could potentially play a role. In the end, we identified 59 potential fcLMR pairs that correlate with a similar set of metabolic pathways and functions in humans and mice (Figure 1*C* and Supplementary Table 3).

### Human and mouse fcLMRs in the liver share similar metabolic functions in vivo

To rigorously test the robustness of our selection strategy, we simultaneously examined the function of four pairs of putative fcLMRs (h/mLMR1-4) (Figure 2*A*). They all met the above-mentioned selection criteria. The dominantly expressed transcripts of some of the lncRNAs defined in our novel annotations were very different from those in the reference annotations (GENCODE v33 and GENCODE vM24). mLMR2 and mLMR3 had no matching transcripts in reference annotations, and the isoforms of other LMRs also showed substantial differences. This indicates that these lncRNA pairs would have never been discovered if current reference annotations were used. Based on the novel annotations, we have successfully cloned all lncRNAs (Supplementary Figure 2A) and performed in vitro translation to validate their noncoding nature (Supplementary Figure 2B). These potential fcLMRs also showed consistent responses to specific metabolic treatments in the chimeric human/mouse livers. For example, h/mLMR2 were upregulated by fasting and h/mLMR4 were both decreased by FXR activation (Figure 2*B*).

**Figure 2.**
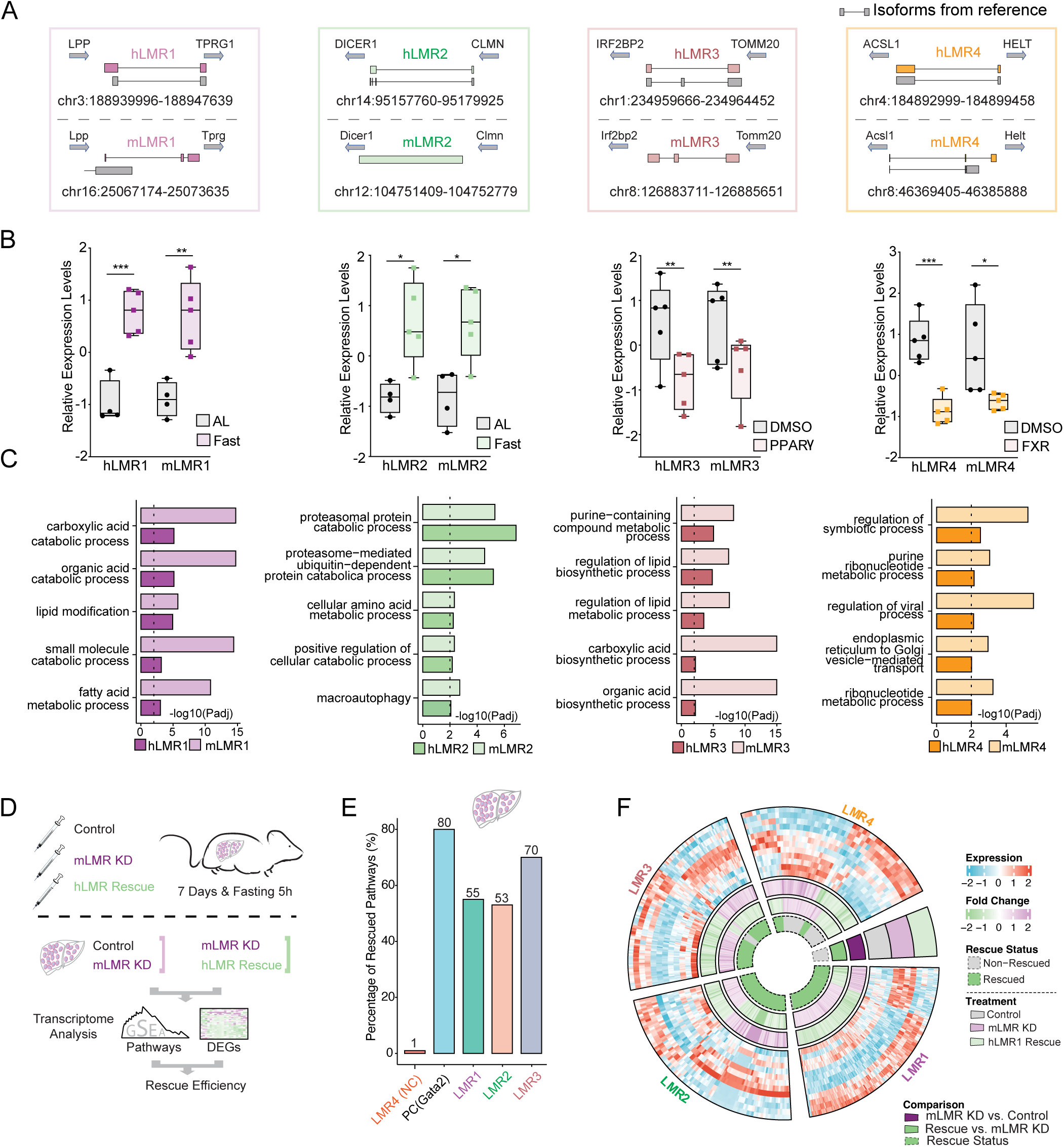
Human and mouse fcLMRs share similar metabolic functions in vivo (A) Diagram of relative positions of human and mouse LMRs with respect to their orthologous genes. Transcripts from reference databases (GENCODE v33 and GENCODE vM24) were shown in grey. For genes with several isoforms, the transcript that is dominantly expressed was displayed. (B) Boxplots of the relative gene expression levels of human and mouse LMRs in humanized livers under different treatment conditions. The gene expression levels were normalized by Z-score across samples. *p<0.05, **p<0.01 and ***p<0.001. (C) Bar plots of the pathways that were commonly enriched in human and mouse LMRs. The length of the bars represented the enrichment significance of the pathways (-log10(adjusted p value)). (D) Schematic diagram of the rescue experiment in mice. Mice were injected with three groups of viruses, including control viruses for knockdown and overexpression (Lac+pAdv5, control group), mLMR knockdown viruses and control viruses for overexpression (mouse-lncRNA KD+pAdv5, mLMR knockdown group, mLMR KD group), and mLMR knockdown viruses and hLMR overexpression group (mouse- lncRNA KD+human-lncRNA OE, hLMR rescue group). The efficiency of rescue was evaluated by comparing the differential pathways and genes between the mLMR knockdown group and control group, as well as between the hLMR rescue group and mLMR KD group. (E) Bar plot of the percentages of rescued pathways for each LMR. (F) Circular diagram of the expression levels of differentially expressed genes in the control group, mLMR knockdown (KD) group, and hLMR rescue group. These differentially expressed genes were identified as commonly and significantly changed genes during the mLMR knockdown and hLMR rescue processes. The fold changes of the genes in the mLMR knockdown and hLMR rescue groups were summarized on the inner circle. The rescue statuses of these genes were also summarized in the center of the diagram, with green representing rescued genes and gray representing non-rescued genes.

Pathway analysis of the correlated genes shows that h/mLMRs in each pair are associated with similar metabolic processes. For instance, both h/mLMR1 are linked to fatty acid metabolism and h/mLMR2 are associated with cellular amino acid metabolism (Figure 2*C*).

Rescue experiments in mice offer a valuable means to verify the similarity of a fcLMR pair’s physiological function. By suppressing mLMRs in mice and observing the impact on gene regulation, followed by overexpressing hLMRs, we can determine the ability of hLMRs to rescue the loss-of-function effects of mLMRs at a physiological level. Should hLMRs exhibit a reversal of the loss-of-function, this would suggest that they have similar in vivo functions to mLMRs. To do this, we suppressed the expression of mLMRs in mouse livers, while overexpressing the corresponding hLMRs as a rescue group (Figure 2*D*). RT-qPCR results showed a minimum 50% reduction in the expression levels of mLMRs in the knockdown groups compared to the controls (Supplementary Figure 2C-F). Furthermore, overexpression (OE) of hLMRs was effective in all cases (Supplementary Figure 2C-F). RNA-seq was performed to analyze the expression levels of the transcriptomes (Figure 2*D*).

Regulation of gene expression is among the most well-established functions of lncRNAs, so we used changes in transcriptome-level gene expression and pathway enrichment as a proxy to evaluate the functional similarity of the selected fcLMR pairs. To verify the feasibility of this approach, we first examined the rescue efficacy of a protein coding gene. We analyzed a public RNA-seq dataset which compared gene expression between hematopoietic progenitor cells from Gata2-deficient mice expressing Gata2 and control (empty vector)^11^ and found that 80% of biological pathways regulated by Gata2 can be rescued (Figure 2*E*). We then used the same approach to evaluate the function of fcLMRs by comparing gene expressions in mLMR knockdown mice with those with concurrent mLMR knockdown and hLMR expression. We found that one hLMR, hLMR4, did not rescue the function of its corresponding mLMR, which we thereafter used as a negative control (Figure 2*E*). It is possible that hLMR4 and mLMR4 do not share similar function or they function in cis, which cannot be evaluated by the current strategy. In contrast, all three remaining hLMRs, hLMR1-3, rescued 55%, 53% and 70% of the pathways regulated by mLMR1-3, respectively (Figure 2*E* and Supplementary Table 4). Consistent with the rescued pathways, we also found that hundreds of genes that were altered by knockdown of mLMR1-3 displayed opposite expression patterns in the rescue groups, which were similar to those in the controls (Figure 2*F* and Supplementary Table 4). These data implicate that 3 out of 4 tested human/mouse LMRs have similar functions, indicating that fcLMRs are common occurrence and our selection strategy can identify them with high efficiency.

### hLMR1 and mLMR1 modulate lipid metabolism

After confirming that three hLMRs could rescue the function of their corresponding mLMRs in the liver based on gene regulation, we next used hLMR1 and mLMR1 as an example to investigate if mLMR knockdown leads to changes in metabolic response and if these changes can be rescued by their corresponding hLMRs. h/mLMR1 are located in syntenic regions flanked by the same orthologous coding genes, LPP/Lpp and TPRG1/Tprg1, in human and mouse genomes. Long-read direct RNA-sequencing reveals that the transcript sequences of these lncRNAs in the liver are different from those in the reference annotations (Figure 2*A*). Correlation analysis suggested that hLMR1 and mLMR1 may play a role in fatty acid metabolism (Figure 2*C*). Additionally, hLMR1 was found to be up-regulated in livers of patients with nonalcoholic fatty liver disease (NAFLD), and mLMR1 was induced by high-fat diet feeding, suggesting a similar regulation of both lncRNAs in liver metabolic disorders (Figure 3*A*).

**Figure 3.**
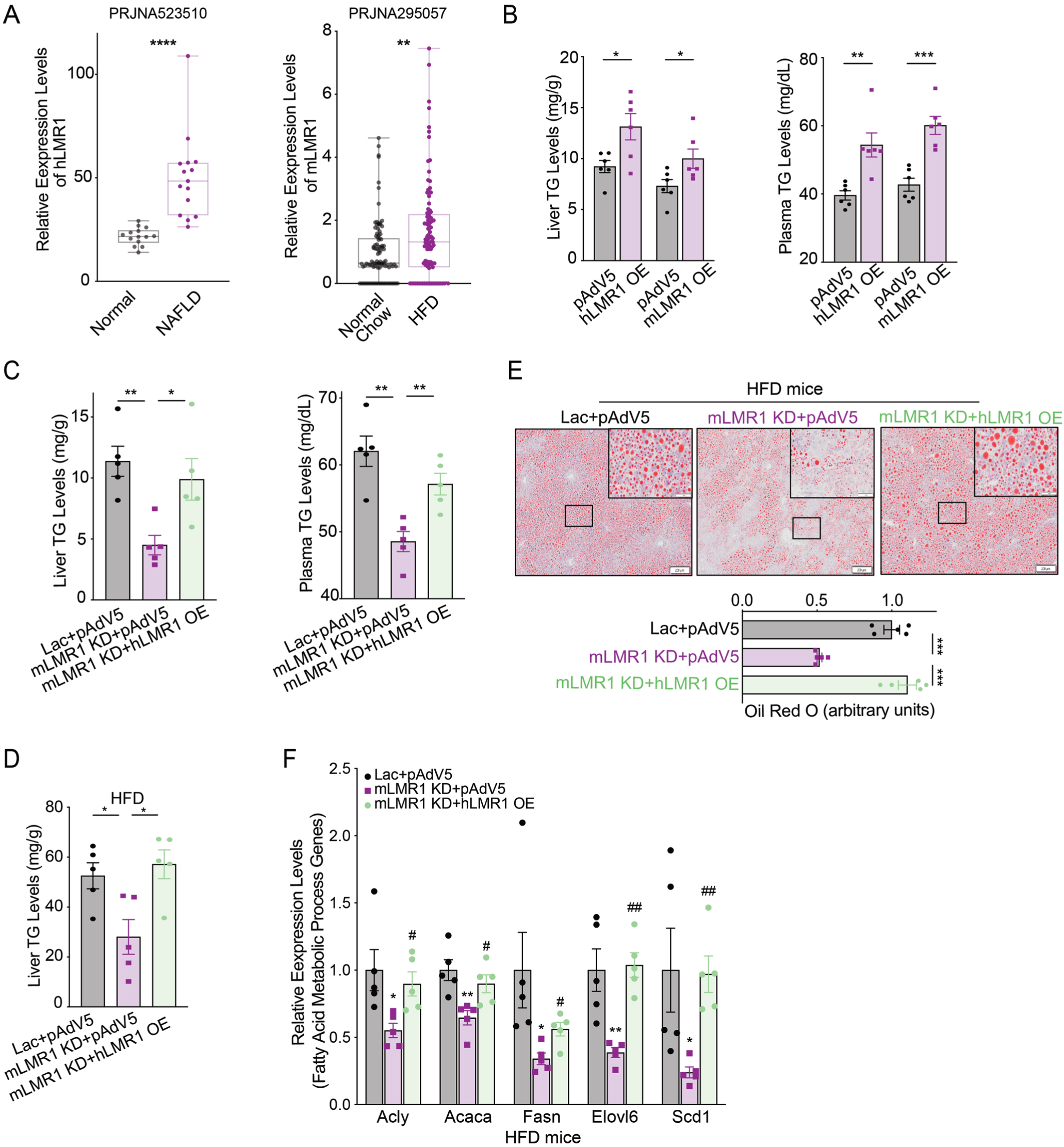
hLMR1 and mLMR1 modulate lipid metabolism (A) Left: the relative expression levels of hLMR1 in the livers of healthy, nonalcoholic fatty liver patients. ****p<0.0001. Right: the relative expression levels of mLMR1 in the livers of normal chow, high fatty diet treated mice. (B) Levels of triglyceride (TG) in liver (left) and plasma (right) in mice with overexpression of hLMR1 or mLMR1 (n=6 per group). (C) Levels of TG in liver (left) and plasma (right) in mice receiving the following adenovirus combinations: Lac+pAdv5 (control group), mLMR1 KD+pAdv5 (mLMR1 knockdown group) or mouse-LMR1 KD+hLMR1 OE (hLMR1 rescue group) (n=5 per group). (D) Levels of TG in liver of high fatty diet (HFD)- fed mice receiving the following adenovirus combinations: Lac+pAdv5, mLMR1 KD+pAdv5, or mLMR1 KD+hLMR1 (n=5 per group, treated with HFD for 9 weeks). (E) The representative images (x2) of oil red staining of liver sections from HFD-fed mice receiving the following adenovirus combinations: Lac+pAdv5, mLMR1 KD+pAdv5, or mLMR1 KD+hLMR1 (Top). (F) The relative expression levels of representative genes from the fatty acid metabolic process pathway in HFD-fed mice receiving the following adenovirus combinations: Lac+pAdv5, mLMR1 KD+pAdv5, or mLMR1 KD+hLMR1 (n=5 per group). Data represent mean ± SEM. *, comparison between mLMR1 KD+pAdv5 and Lac+pAdv5. ^#^, comparison between mLMR1 KD+hLMR1 OE and mLMR1 KD+pAdv5. Data represent mean ± SEM, * or ^#^ p<0.05, ** or ^##^ p<0.01, *** or ^###^ p<0.001.

To evaluate the role of h/mLMR1 in lipid metabolism, we overexpressed them in mouse livers (Supplementary Figure 3A and B) and found that both increased triglyceride (TG) levels in the liver and plasma (Figure 3*B*). Consistently, overexpression of hLMR1 increased the protein levels of Scd1, Fasn, and Acaca, key enzymes in fatty acid pathways (Supplementary Figure 3C). Conversely, we found that mLMR1 knockdown led to lower TG levels in livers and plasma, and these phenotypes were fully rescued by concurrent expression of hLMR1 (Figure 3*C*). To understand the molecular basis of the changes in TG levels, we examined the expression of key genes in the fatty acid synthesis pathway and found that their expression levels were decreased in the livers of the KD group and recovered in the rescue group, a pattern of regulation similar to that of the TG levels (Supplementary Figure 3D). These results suggest that h/mLMR1 might regulate TG levels by modulating the expression of lipogenic genes.

To explore the potential role of hLMR1 and mLMR1 in liver disorders, we conducted a rescue assay in mice fed with a high-fat diet, which is known to induce hepatic steatosis (Supplementary Figure 3E and F). Consistent with the results in normal chow-fed mice, the mLMR1 KD group displayed lower liver TG levels compared to those in the control group, which were also able to be rescued by expression of hLMR1 (Figure 3*D*). These results were further supported by oil red staining, which showed that the mLMR1 KD group had fewer lipid droplets than the control group did, while concurrent hLMR1 expression reversed this pattern (Figure 3*E*). Additionally, genes involved in the fatty acid synthesis such as Acly, Acaca, Fasn, Elovl6, and Scd1 were all significantly downregulated in the mLMR1 KD group and rescued to levels comparable to those in controls by hLMR1 expression (Figure 3*F*). These results indicate that hLMR1 and mLMR1 may play a similar role in inducing steatosis, with dysregulated hLMR1 in patients with NAFLD potentially being pathogenic.

### Both hLMR1 and mLMR1 bind to PABPC1 and inhibit protein translation

We explored the molecular mechanism that h/mLMR1 regulate the expression of lipogenic genes. It is known that lncRNAs can act in cis to regulate the expression of their neighboring genes or in trans via interactions with proteins, RNAs or DNAs. Our overexpression and rescue experiments in Figure 3 indicate that h/mLMR1 can function in trans. Consistently, we found that the expression levels of Lpp and Tprg1, the neighboring genes of mLMR1, were not changed by mLMR1 knockdown or hLMR1 overexpression (data not shown). Next, we performed an RNA pull-down using biotinylated hLMR1 followed by mass spectrometry analysis to identify hLMR1- protein binding interactions. This approach led us to identify Poly(A)-binding protein C1 (PABPC1) as a binding partner of hLMR1, which we confirmed by anti-PABPC1 immunoblotting (Figure 4*A*). Intriguingly, we found that mLMR1 also interacted with PABPC1 (Figure 4*A*). Of note, PABPC1 is known to be able to bind to the poly(A) tails of RNAs^12^ but its binding to h/mLMR1 is not dependent on poly(A) tails, as hLMR1 and mLMR1 used in our pull-down experiments are devoid of these sequences. Furthermore, RNA Immunoprecipitation analysis of humanized liver samples showed that immunoprecipitation of PABPC1 enriched hLMR1 and mLMR1 by more than 200- and 100-fold, respectively, relative to the controls (IgG) (Figure 4*B*). PABPC1 is known to play a crucial role in the regulation of mRNA translation^12^. Indeed, our analysis of publicly available liver cancer databases (The Cancer Genome Atlas) revealed that the group of samples with high Pabpc1 expression levels exhibited higher overall protein abundance (Figure 4*C*), supporting a positive correlation between PABPC1 expression and protein translation in the liver. To investigate the function of PABPC1 in the liver, we overexpressed and knocked down mouse Pabpc1 in the liver (Supplementary Figure 4A and 4B) and evaluated protein translation by performing a polysome fractionation assay. This analysis revealed that Pabpc1 overexpression led to a significant increase in the polysome ratio across all fractions (Figure 4*D*), indicating higher protein translation rates in the liver. Conversely, Pabpc1 knockdown resulted in reduced protein translation activity (Figure 4*E*), consistent with the reported role of Pabpc1 in protein translation.

**Figure 4.**
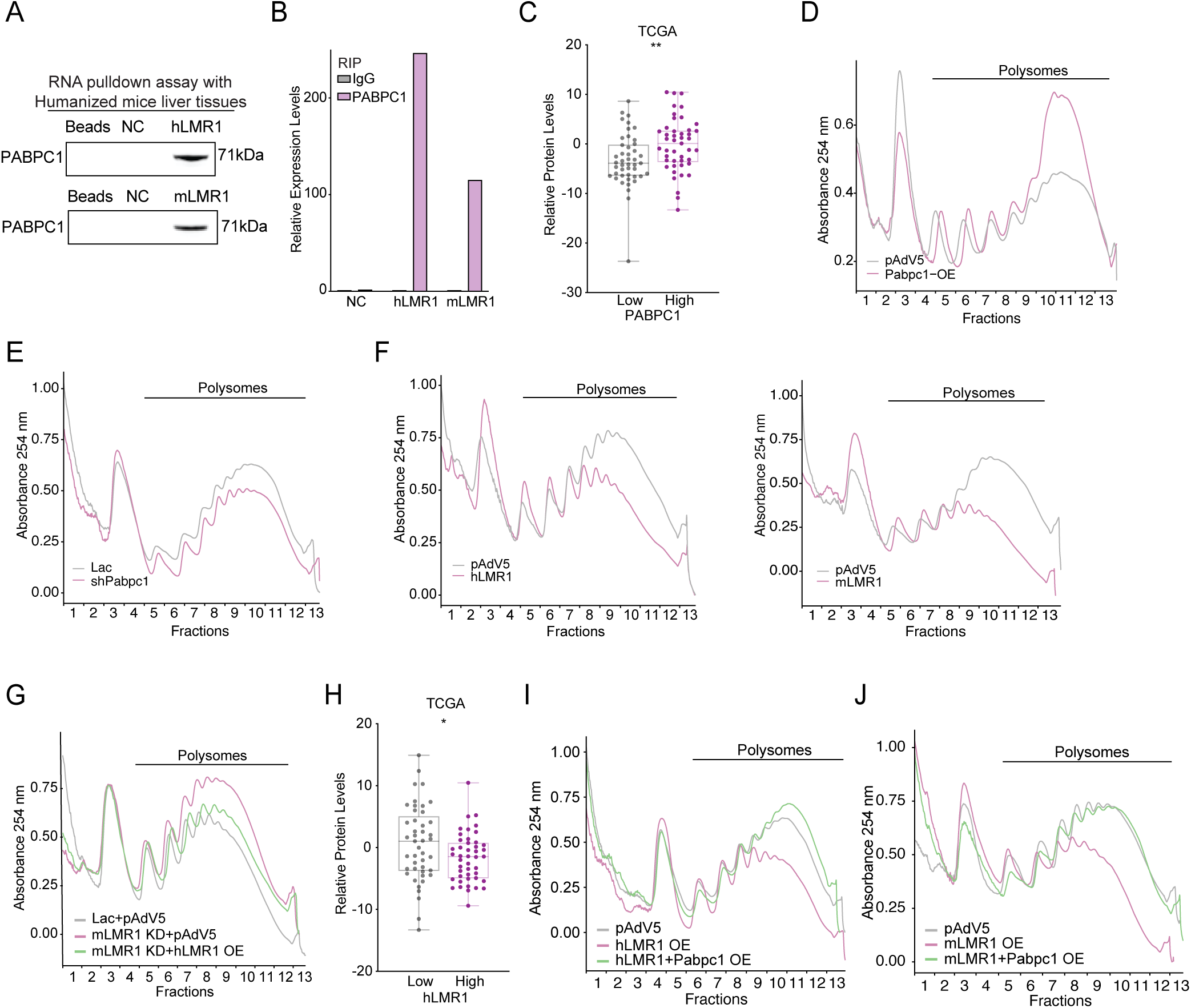
Both hLMR1 and mLMR1 bind to PABPC1 and inhibit protein translation (A) The RNA pull-down results of hLMR1 and mLMR1 in the pooled humanized mouse livers. The Beads group represented beads-only pull-down, while the NC group represented anti-sense sequence pull-down for hLMR1 and mLMR1, respectively. (B) The relative expression levels of NC, hLMR1, and mLMR1 from RNA immunoprecipitation (RIP) in the humanized liver. Human LINC02499 was used as a negative control (NC). (C) Box plot of the relative expression levels of total proteins in the TCGA database liver tissue samples (TCGA, LIHC) with high and low PABPC1 expression. **p<0.01. (D and E) Sucrose gradient absorbance profiles of mouse livers with overexpression (D) or knockdown of Pabpc1 (E). Liver samples from the same treatment group were pooled together, and the polysome fractions were labeled. (F) Sucrose gradient absorbance profiles of mouse livers with hLMR1 (Left) or mLMR1 overexpression (Right). (G) Sucrose gradient absorbance profiles of livers from mouse receiving the following adenovirus combinations: Lac+pAdv5, mLMR1 KD+pAdv5, or mLMR1 KD+hLMR1 OE. (H) Box plot of the relative expression levels of total proteins in the TCGA database liver tissue samples (TCGA, LIHC) with high (n=45) and low (n=45) hLMR1 expression. *p<0.05. (I) Sucrose gradient absorbance profiles were obtained from pooled liver samples of mice receiving the following adenovirus combinations: pAdv5 (control group), hLMR1 overexpression (hLMR1 OE), and co- overexpression of h LMR1 and Pabpc1 (hLMR1+Pabpc1 OE). (J) Sucrose gradient absorbance profiles were obtained from pooled liver samples of mice receiving the following adenovirus combinations: pAdv5 (control group), mLMR1 overexpression (mLMR1 OE), and co- overexpression of m LMR1 and Pabpc1 (mLMR1+Pabpc1 OE).

Given the binding of Pabpc1 to LMR1, we examined if LMR1 also plays a role in regulating protein translation. Polysome fractionation of livers overexpressing hLMR1 or mLMR1 showed that both displayed a significant reduction in polysome abundance (Figure 4*F*). Conversely, mLMR1 knockdown increased polysome abundance, and critically this increase could be rescued by concurrent expression of hLMR1 (Figure 4*G*). These data collectively support that both hLMR1 and mLMR1 have a similar role in suppressing protein translation. Furthermore, increased expression levels of hLMR1 in human liver tissues were associated with decreased protein abundance (Figure 4*H*), again supporting hLMR1’s role in regulating protein translation in humans. As both hLMR1 and mLMR1 specifically bind to PABPC1, they might suppress protein translation via inhibiting the function of PABPC1. To investigate this possibility, we overexpressed Pabpc1 in hLMR1- or mLMR1-expressing mouse livers and found that it rescued the decreased translation in both cases (Figure 4*I* and J, Supplementary Figure 4C-F), indicating that PABPC1 is a downstream effector of h/mLMR1 in regulating protein translation.

### Inhibition of protein translation by h/LMR1-PABPC1 complex led to increased lipogenesis

Protein translation and lipid synthesis are interconnected and can affect each other in various ways^13^. Given the inhibitory role of h/mLMR1 on PABPC1 and protein translation, we explored if these lncRNAs regulate lipid metabolism via PABPC1—potentially by modulating translation rate. We have shown that overexpression of either hLMR1 or mLMR1 increases plasma TG levels. Intriguingly, when we overexpressed Pabpc1 in hLMR1 or mLMR1 OE mice, the elevated levels of plasma TG were effectively reversed(Figure 5*A* and *B*, Supplementary Figure 4C-F). Moreover,

**Figure 5.**
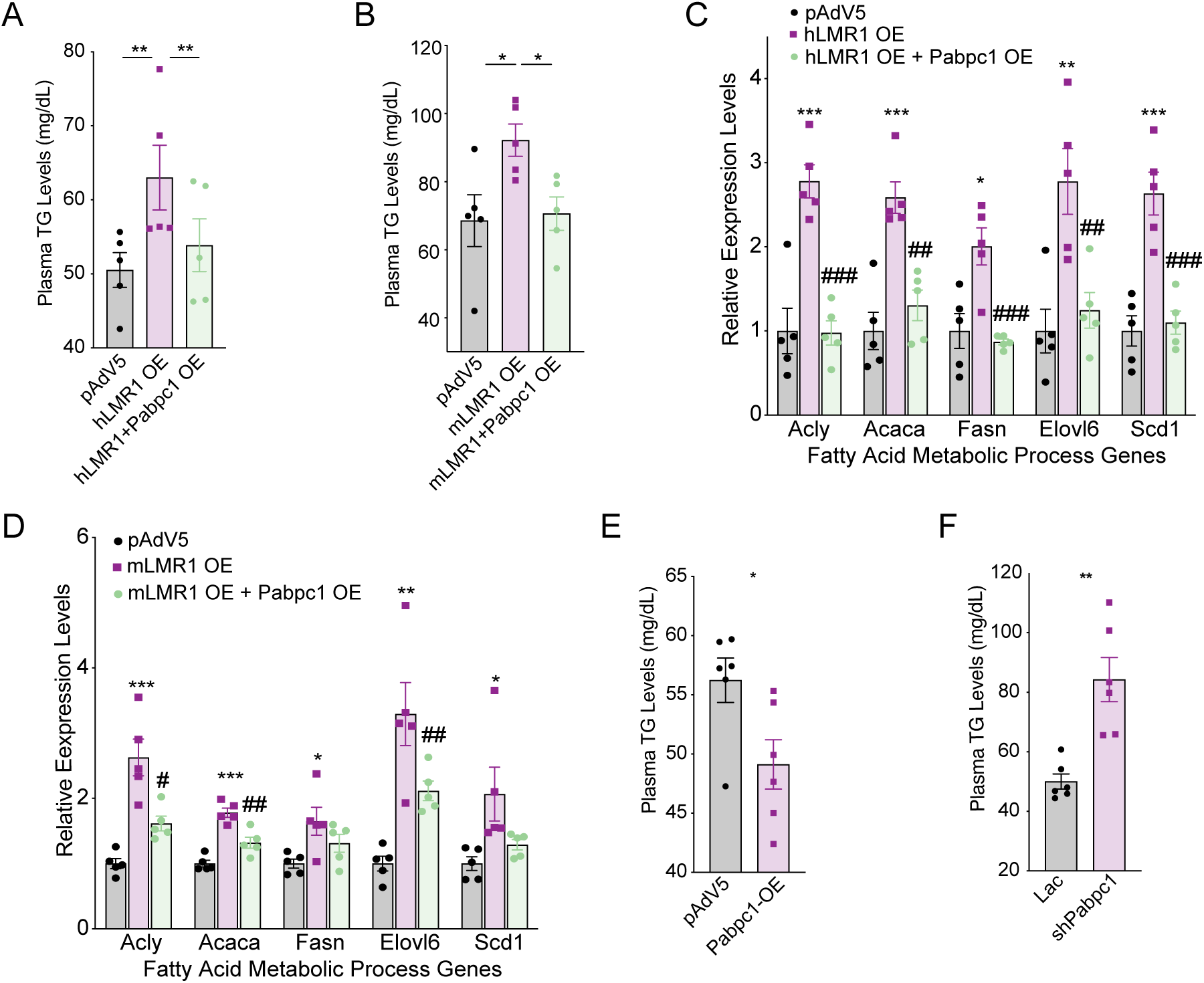
Inhibition of protein translation by h/mLMR1-PABPC1 complex led to increased lipogenesis (A) The levels of plasma TG in mice receiving the following adenovirus combinations: pAdv5 (control group), hLMR1 overexpression (hLMR1 OE), and co-overexpression of hLMR1 and Pabpc1 (hLMR1+Pabpc1 OE). (B) The levels of plasma TG in mice receiving the following adenovirus combinations: pAdv5 (control group), mLMR1 overexpression (mLMR1 OE), and co- overexpression of mLMR1 and Pabpc1 (mLMR1+Pabpc1 OE). (C) The relative expression levels of representative genes from the fatty acid metabolic process pathway were measured in livers of mice receiving the following adenovirus combinations: pAdv5, hLMR1 OE, and hLMR1+Pabpc1 OE. *, comparison between h/mLMR1 OE and pAdv5. ^#^, comparison between hLMR1+Pabpc1 OE and hLMR1 OE. (D) The relative expression levels of representative genes from the fatty acid metabolic process pathway were measured in livers of mice receiving the following adenovirus combinations: pAdv5, mLMR1 OE, and mLMR1+Pabpc1 OE. *, comparison between h/mLMR1 OE and pAdv5. ^#^, comparison between mLMR1+Pabpc1 OE and mLMR1 OE. (E) The levels of plasma TG in mice receiving pAdv5 (control group) or Pabpc1 overexpression (Pabpc1 OE) adenoviruses (n=6 per group). (F) The levels of plasma TG in mice receiving pAdv5 (control group) or Pabpc1 knockdown (shPabpc1) adenoviruses (n=6 per group). Data represent mean ± SEM, * or ^#^ p<0.05, ** or ^##^ p<0.01, *** or ^###^ p<0.001.

Pabpc1 expression also mitigates the increased expression levels of lipogenic genes induced by hLMR1 or mLMR1 overexpression (Figure 5*C* and *D*). To further confirm the role of PABPC1 in lipid metabolism, we overexpressed Pabpc1 in mouse livers and found that it led to a significant decrease in plasma TG levels, while Pabpc1 knockdown had the opposite effect (Figure 5*E* and *F*). These findings suggest that h/mLMR1 might indirectly alter lipid metabolism by regulating protein translation via binding to Pabpc1.

### h/LMR1 regulate lipid metabolism via an amino acids-mTOR- SREBP1 axis

Next, we explored how alterations in PABPC1 function and protein translation might modulate lipid synthesis in the liver. Macronutrients, i.e., proteins, carbohydrates, and fats are inter- convertible in the body, and the levels of these metabolites are important signals that regulate the overall metabolic flux. For example, elevated levels of amino acids and glucose can both induce the expression of lipogenic genes by activating mTOR and chREBP, respectively^14, 15^. As h/mLMR1 inhibits PABPC1 and protein translation, it might affect the levels of amino acids and subsequently modulate lipogenesis via mTOR. Indeed, the levels of amino acids in the livers were significantly increased by either hLMR1 or mLMR1 overexpression (Supplementary Figure 5A and B) as well as PABPC1 knockdown (Supplementary Figure 5C). More critically, the levels of amino acids were significantly reduced by mLMR1 knockdown and were rescued by concurrent expression of hLMR1 (Figure 6*A*). Furthermore, the increased amino acid levels by hLMR1 or mLMR1 could be reversed by concurrent expression of PABPC1 (Figure 6*B* and *C*) further supporting that these lncRNAs inhibit protein translation via PABPC1. It is well established that amino acids can activate mTOR, a strong driver of lipogenesis that acts on lipin1 or S6K to increase the nuclear accumulation of SREBPs and expression of lipogenic genes^15^. We found that mTOR phosphorylation was decreased by mLMR1 knockdown and was rescued by concurrent expression of hLMR1 (Figure 6*D*). Conversely, mTOR phosphorylation was increased by hLMR1 or mLMR1 overexpression and reversed by concurrent expression of PABPC1 (Figure 6*E* and *F*). Finally, we examined the SREBP1 transcriptional activities towards lipogenic genes. An anti-SREBP1 chromatin immunoprecipitation assay revealed that SREBP1 binding to promoters/enhancers of lipogenic genes fully mirrored mTOR phosphorylation in all abovementioned conditions, i.e, decreased by mLMR1 knockdown and rescued by concurrent expression of hLMR1 (Figure 6*G*); and increased by hLMR1 or mLMR1 overexpression and reversed by concurrent expression of PABPC1 (Figure 6*H* and *I*). Taken together, our results here support that h/mLMR1 binds to PABPC1 and inhibits translation, which subsequently activates an amino acid-mTOR -SREBP1 axis to increase lipogenic gene expression.

**Figure 6.**
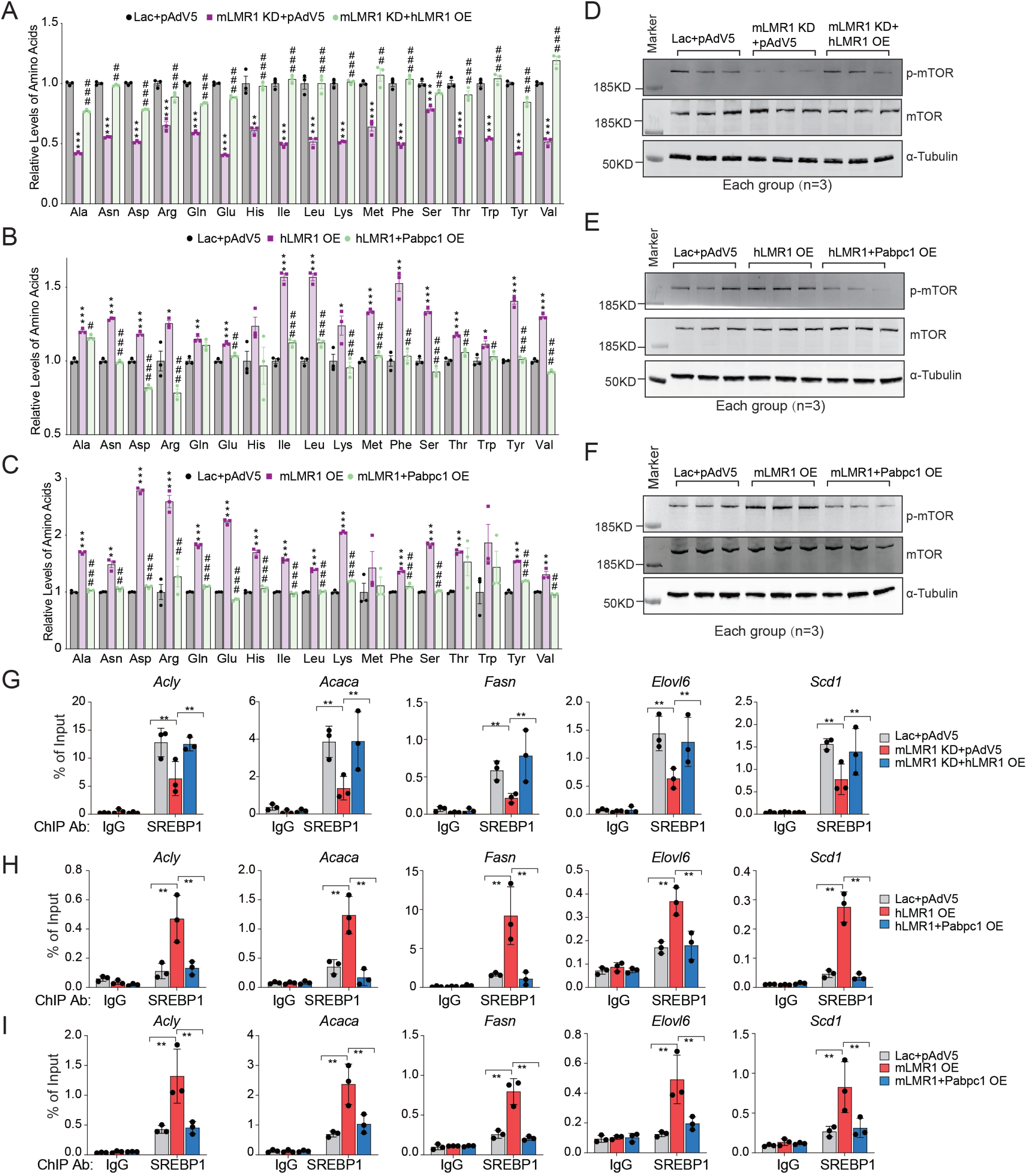
h/mLMR1 regulate lipid metabolism via an amino acids-mTOR-SREBP1 axis (A) Relative levels of amino acids in the livers of mice fed a high-fat diet (HFD) and receiving one of the following adenovirus combinations: Lac+pAdv5 (Ctrl), mLMR1 KD+pAdv5 (mLMR1 KD), or mLMR1 KD+hLMR1 (mice treated with HFD for 9 weeks). *, comparison between mLMR1 KD and Ctrl. ^#^, comparison between mLMR1 KD+hLMR1 and mLMR1 KD. (B) The relative levels of amino acids were measured in the livers of mice receiving the following adenovirus combinations: pAdv5 (V5), hLMR1 OE, and h LMR1+Pabpc1 OE. *, comparison between hLMR1 OE and V5. ^#^, comparison between hLMR1+Pabpc1 OE and hLMR1 OE. (C) The relative levels of amino acids were measured in the livers of mice receiving the following adenovirus combinations: pAdv5 (V5), mLMR1 OE, and mLMR1+Pabpc1 OE. *, comparison between mLMR1 OE and V5. ^#^, comparison between mLMR1+Pabpc1 OE and mLMR1 OE. (D- F) Immunoblot analyses of phosphor-mTOR (p-mTOR), total mTOR and α-Tubulin proteins in mice subjected to different treatments (n=3 per group): (D) Lac+pAdv5 (Ctrl), mLMR1 KD+pAdv5 (mLMR1 KD), or mLMR1 KD+hLMR1 OE; (E) pAdv5 (V5), hLMR1 OE, or hLMR1+Pabpc1 OE; (F) pAdv5 (V5), mLMR1 OE, or mLMR1 and Pabpc1 OE. (G-I) Anti- SREBP1 and IgG chromatin immunoprecipitation (ChIP) analyses of livers in mice subjected to different treatments (n=3 per group): (G) Lac+pAdv5 (Ctrl), mLMR1 KD+pAdv5 (mLMR1 KD), or mLMR1 KD+hLMR1 OE; (H) pAdv5 (V5), hLMR1 OE, or hLMR1+Pabpc1 OE; (I) pAdv5 (V5), mLMR1 OE, or mLMR1 and Pabpc1 OE. Data represent mean ± SEM, * or ^#^ p<0.05, ** or ^##^ p<0.01, *** or ^###^ p<0.001.

### Short motifs of similar structure on h/mLMR1 bind to PABPC1 and share similar in vivo function

To understand the molecular basis of the interaction between h/mLMR1 and PABPC1, we generated a series of truncated human PABPC1 constructs with deletions of individual functional domains (Supplementary Figure 6A and B) and evaluated their binding affinity to h/mLMR1. Our results showed that both MLLE and RRM2 domains of PABPC1 were crucial for binding to h/mLMR1 (Figure 7*A*). More importantly, for both h/mLMR1, the reduced binding affinities towards each truncated PABPC1 were similar (Figure 7*A*), suggesting that they might bind to the same domain on PABPC1. To further investigate this possibility, we performed a binding competition assay and found that h/mLMR1 could effectively compete with each other’s binding to PABPC1 (Figure 7*B* and Supplementary Figure 6C), once again supporting the idea that they might share the same binding site on PABPC1.

**Figure 7.**
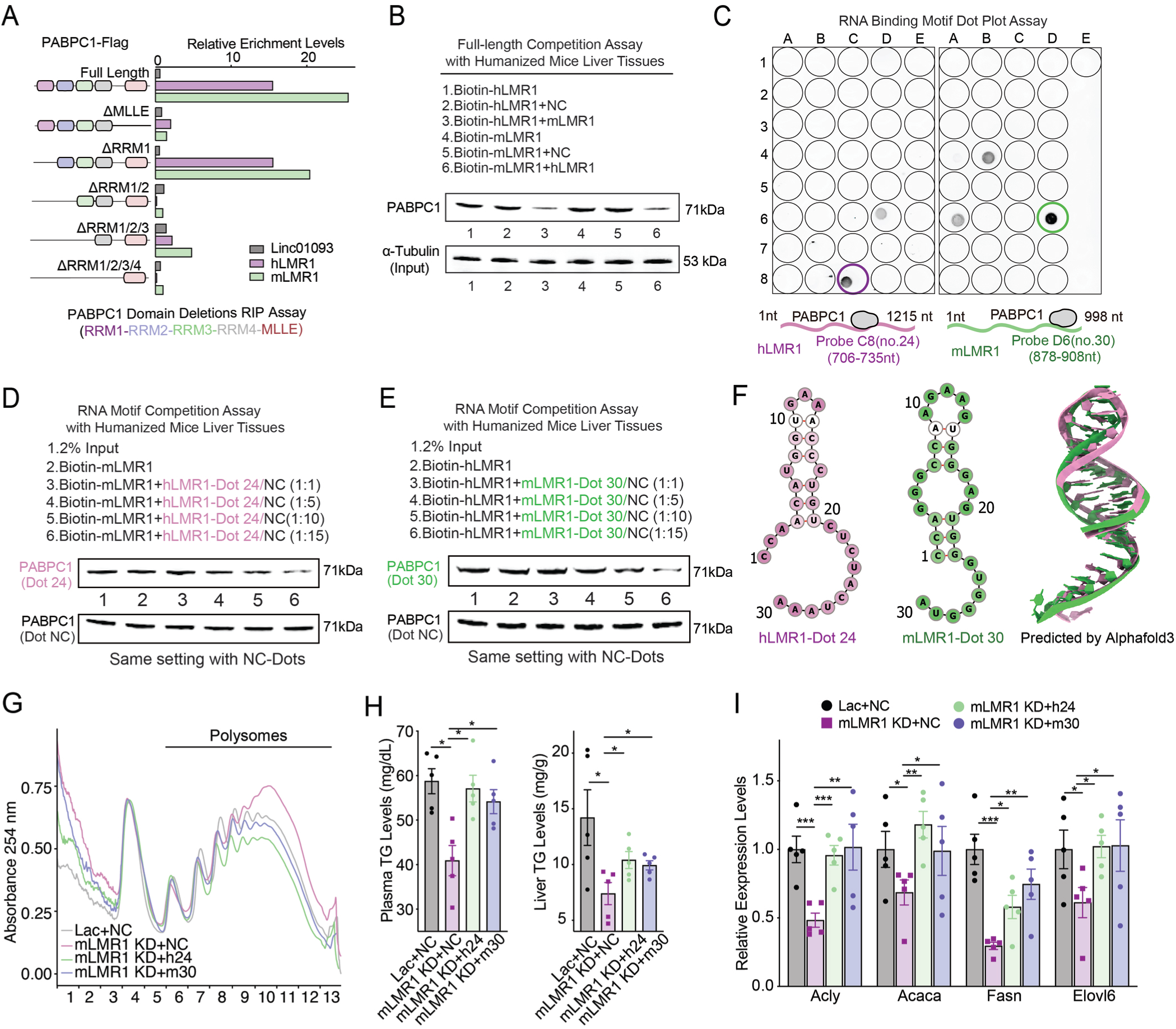
Short motifs of similar structure on h/mLMR1 bind to PABPC1 and share similar in vivo function (A) The enrichment levels of h/mLMR1 in immunoprecipitation (RIP) assay using Flag-tagged PABPC1 isoforms shown on the left. (B) The competition assay between hLMR1 and mLMR1 for binding to PABPC1. The bound PABPC1 were detected by immunoblotting using an anti-Flag antibody. The experimental conditions for different groups were annotated above the image. (C) In vitro RNA-protein binding determined by dot blot assay using in vitro–transcribed biotinylated hLMR1 (left) and mLMR1 (right). (D and E) The competition assay between fragment #24 of hLMR1 (D) and fragment #30 of mLMR1 (E) for binding to PABPC1. Western blotting analysis showing the effect of competition between LMR1 fragments and full-length LMR1 for binding to PABPC1. The experimental conditions for different groups were annotated above the image. (F) Predicted secondary and tertiary structures of hLMR1 fragment 24 (hLMR1 #24) and mLMR1 fragment 30 (mLMR1 #30). The secondary structures were predicted using RNAfold, and the tertiary structures were predicted using Alphafold3. (G) Polysome analysis of liver samples from mice receiving the following adenovirus combinations: Lac+NC (knockdown control and LMR1 fragment overexpression control, control group), mLMR1 KD+NC (mLMR1 knock down and LMR1 fragment overexpression control, mLMR1 knockdown group), mLMR1 KD+h24 (mLMR1 knockdown and hLMR1 fragment #24 overexpression, hLMR1 fragment #24 rescue group), and mLMR1 KD+m30 (mLMR1 knockdown and mLMR1 fragment #30 overexpression, mLMR1 fragment #30 rescue group). (H) The levels of triglyceride in plasma and liver in mice shown in (G). (I) The relative expression levels of representative genes in the fatty acid metabolic process pathway were measured in mouse livers described in (G). Data represent mean ± SEM. *p<0.05, **p<0.01, ***p<0.001.

To identify the specific regions on h/mLMR1 that mediate their PABPC1 binding, we performed a dot blot assay to scan each lncRNA at a 30-nt interval to map their binding motifs. This experiment showed that region #24 (CCAACAUGGUGAAACCCUGUCUCUACUAAA, C8 well) on hLMR1 and region #30 (CCAGGGCCAAGAAGUGGGAGUGGGUGGGUA, D6 well) on mLMR1 exhibited the strongest binding with PABPC1 (Figure 7*C* and Supplementary Figure 6 *D*). To determine the significance of #24 and #30 to the interaction between h/mLMR1 and PABPC1, we performed competitive binding experiments using both hLMR1 (#24) and mLMR1 (#30) with full-length h/mLMR1. Results from this assay showed that hLMR1 (#24) was able to effectively compete with full-length mLMR1 for its binding to PABPC1, and this competition became more pronounced with increasing doses (Figure 7*D*, Supplementary Figure 6 *E*). A similar observation was made for mLMR1 (#30), which robustly competed with full-length hLMR1 for its binding to PABPC1 (Figure 7*E*). These findings indicate that hLMR1 (#24) and mLMR1 (#30) play a crucial role in the interaction between LMRs and PABPC1. Furthermore, the predicted RNA secondary structures and tertiary structures of hLMR1 (#24) and mLMR1 (#30) exhibited a high degree of similarity (Figure 7*F*) which could be the underlying reason that they both interact with a specific domain of PABPC1.

Finally, to determine the functional role of hLMR1 (#24) and mLMR1 (#30) in vivo, we reintroduced them into mLMR1 knockdown mice (Supplementary Figure 6F) and found that either motif could reverse the increased polysome abundance (Figure 7*G*). Intriguingly, both motifs also partially restored the decreased TG levels in plasma and liver (Figure 7*H*). Moreover, the decreased expression levels of genes in fatty acid metabolic processes were also rescued in hLMR1 (#24) and mLMR1 (#30) rescue groups, and the patterns of regulation were similar to those of TG levels (Figure 7*I*). These findings suggest that the specific motifs on hLMR1 and mLMR1 are essential to their role as regulators of fatty acid metabolism, and their functional similarity might be mediated by their interaction with PABPC1 through a conserved docking mechanism.

## Discussion

In summary, our work supports the concept of functionally conserved lncRNA metabolic regulators (fcLMRs) and introduces an innovative approach to exploring the physiological role of human liver lncRNAs in mouse livers. We established a function-based pipeline to systematically identify potential fcLMRs in human and mouse livers, confirming the legitimacy of 3 out of 4 fcLMRs through in vivo rescue experiments in mice. The high success rate in defining fcLMRs can be attributed to the robustness of our selection strategy. We curated accurate annotations of human and mouse liver lncRNAs, enabling the detection of syntenic lncRNAs absent in current reference annotations. Additionally, we determined the precise regulation of human and mouse lncRNAs under identical in vivo metabolic conditions using liver-specific humanized mice and conducted a correlation analysis of lncRNAs based on gene expression in humanized livers with an identical genetic background and housing environment. Notably, the expression levels of three characterized human LMRs (hLMRs) were significantly altered in the livers of patients with nonalcoholic fatty liver disease (NAFLD) (Supplementary Figure 7A), suggesting a potential role in the disorder. Our in-depth analysis of h/mLMR1 revealed that fcLMRs can regulate metabolic fluxes by fully engaging with essential protein-coding metabolic regulators such as SREBP1 and mTOR through structurally conserved motifs, which themselves could serve as potential therapeutic targets.

Our results support that fcLMRs in the human liver might be a common occurrence. The number of putative LMR pairs (59) we have identified so far appears to be moderate, potentially due to the extremely stringent selection criteria which could underestimate their real number. In addition, the number of metabolic conditions used to identify similarly regulated lncRNAs can also substantially affect the total quantity of identified fcLMRs. In our current selection, only five metabolic conditions were used, which translated to approximately a 10% increase in the number of similarly regulated lncRNA pairs per condition (Supplementary Figure 7B). Our current rate of discovery (59 out of 2419) is 2.4%. If 5 to 10 additional treatments are utilized, the percentage of putative fcLMRs could potentially reach 5 to 10% or even more. Considering that lncRNAs are highly tissue-specific and many more lncRNAs are expressed in lncRNA-rich organs beyond the liver (Supplementary Figure 7C), the total number of human lncRNAs is almost certainly much larger than those specific to the liver. For example, if the total number of human lncRNAs (100,000) in one of the most inclusive lncRNA databases is used^2^, the estimated number of putative fcLMR pairs can reach 5,000 to 10,000, which is substantial considering that there are only 20,000 conserved protein-coding genes in the human genome. In support of our assessment, lncRNAs that share characteristics of functional conservation in humans and mice can be found in the literature^16,17^. While preparing our manuscript, a new study exploring a similar concept in the context of cell proliferation was published^18^. This additional evidence not only reinforces our argument but also underscores the critical importance of studying fcLMRs, contributing to a deeper understanding of human liver pathophysiology.

Studying fcLMRs in the liver could substantially increase the understanding of metabolic regulation in the human liver and provide novel therapeutic opportunities for human liver disorders. Although genetic evidence has connected an increasing number of human liver lncRNAs to the pathogenesis of human diseases, most of them have not been analyzed in a physiological context, and the lack of a system to study their physiological function has clearly become a major obstacle to unlocking their potential in understanding and treating human liver diseases. To address this challenge, we have attempted to use a liver-specific humanized mouse model to study the in vivo function of hLMRs^7, 19^. Although this model allowed us to establish proof-of-concept examples that non-conserved human lncRNAs have important metabolic roles, it also suffers from several limitations and cannot be used to sufficiently define the pathophysiological role of non-conserved LMRs for clinical translation. It is based on immune-deficient mice, and only hepatocytes in the liver are humanized^20^. More critically, genetic manipulations of human lncRNAs can only be performed on mature hepatocytes, precluding the examination of any developmental impacts of human lncRNAs —often an important consideration for therapy development. All of these problems would be immediately solved if a corresponding mLMR for a hLMR could be defined and studied in conventional mice using the full range of current genetic tools. Considering the projected number of hLMRs, studying their mLMRs could substantially expand the power of human genetics. Conversely, the function of mLMRs defined in mouse genetic studies can also be utilized to understand the pathogenic role of hLMRs, re-establishing the bi-directional information flow between humans and mice that is “broken” by the limited conservation of lncRNAs. Furthermore, our results support that defining functional motifs on fcLMRs can thus serve as a strategy to pinpoint specific drug targets to develop small molecule inhibitors of lncRNA function or lncRNA mimics that have therapeutic effects.

Collectively, our study underscores the prevalence of fcLMRs in the human liver, presenting a promising avenue to significantly advance our comprehension of human liver physiology and metabolic diseases. Employing deep phenotyping of genetic mouse models of mLMRs provides a novel strategy for elucidating the physiological role of human liver lncRNAs. Furthermore, the identification of fcLMR motifs holds the potential to unveil direct therapeutic targets for metabolic disorders. We hope that capitalizing on these exciting possibilities will accelerate our understanding of the human liver physiology and metabolic diseases and the pace at which we target human lncRNAs for the development of effective therapies for huma liver disorders.

## Materials and Methods

### RNA dot-blot assay

Biotinylated full-length RNAs of hLMR1 and mLMR1 were incubated with recombinant myc-tagged PABPC1 proteins in the RNA-protein binding buffer on ice for one hour. Then, the binding mixture was transferred into a pre-cooled 96-well round-bottomed plate and placed on ice with an ultraviolet (UV) crosslinker, and 150 mJ/cm2 radiation was applied for 10 minutes. After UV-crosslinking, the RNAs were partially digested by RNase A for 30 minutes at 37°C. Then, RNA-protein complexes were purified by c-myc tag magnetic beads, and were then treated with proteinase K. The 5-10 µL of biotin-RNA samples were dropped onto nitrocellulose membrane for each well, and incubated with DNA probes at 65°C for 2 hours. Then, the hybridized membrane was washed with washing buffer at 37°C, 50°C, and 65°C for 20 minutes each. Finally, the protein-bound RNA sequence was visualized. Additional details regarding this procedure can be found in the Supplementary Materials and Methods.

### Competition RNA pulldown assay

For the full-length RNA competition assay of hLMR1 and mLMR1, equimolar amounts of biotin-labeled mLMR1 and non-biotin labeled hLMR1 (or biotin-labeled hLMR1 and non-biotin labeled mLMR1) were incubated with precleared humanized mouse liver lysate, following the same procedure used for the RNA pulldown assay. For the hLMR1-dot24 RNA competition with full length mLMR1 or the mLMR1-dot30 RNA competition with full length hLMR1, we prepared four concentration gradients based on the number of moles and incubated them with precleared humanized mouse liver lysate. The subsequent steps were the same as described for the RNA pulldown assay. The samples were then subjected to electrophoresis and immunoblotting with anti-PABPC1 antibody to detect the signals. Additional details regarding this procedure can be found in the Supplementary Materials and Methods.

## Acknowledgments

We thank Yan Luo, Poching Liu and Yuesheng Li (NHLBI DNA Sequencing and Genomics Core) for RNA-seq analysis and Duck-Yeon Lee (NHLBI Biochemistry Core) for free amino acid analysis. The authors gratefully acknowledge the technical assistance of Ms. Megumi Nishiwaki, Mr. Takaya Homma, and Hiroaki Kato.

## Author contributions

CJ, ZL, PL and HC designed the workflow. CJ, ZL, YM, SS, SKP, and SM carried out experiments to analyze all mLMRs and hLMRs. CJ performed all bioinformatics analyses. NY, KK, SU, YO and HS prepared and treated humanized mice. JZ assisted the analysis of the second structures of mLMR1 and hLMR1 and accessed their similarity. CJ, ZL, PL and HC wrote the manuscript. HC conceived and supervised the study.

## Data availability

The raw sequencing data for humanized mice are available at GEO via the SuperSeries dataset GSE224281, which includes multiple SubSeries such as humanized mice RNA-Seq data for dietary treatments (GSE130525) and transcriptional factor agonist treatments (GSE224279), as well as humanized mice nanopore direct RNA sequencing data (GSE224278). The rescue RNA- Seq data are available at GEO with accession number as GSE234310. All data generated or analyzed during this study are included in the article and Supplementary Materials. Data, analytical methods, and research materials in this article will be available to other researchers from the corresponding authors upon reasonable request.

