## Supplementary Materials and Methods for "Systemic identification of functionally conserved lncRNA metabolic regulators in human and mouse livers"

### Preparation and treatments of humanized mice

All animal experiments were conducted in compliance with the guidelines approved by the NHLBI Animal Care and Use Committee or the Animal Care Committee of the Central Institute for Experimental Animals (CIEA, Kanagawa, Japan). TK-NOG mice (Taconic Biosciences, Cat. 12907) aged 8-10 weeks received an intraperitoneal (i.p.) injection of Ganciclovir (APP Pharmaceuticals, NDC: 63323-315-10) at a dose of 20 mg/kg (4µl/g) one week before surgery. Thawed primary human hepatocytes (company) were recovered in Cryopreserved Hepatocyte Recovery Medium (Thermo Fisher Scientific, Cat. CM7000) and resuspended in ice-cold HBSS prior to surgery. Mice were anesthetized with Ketamine/Xylazine by i.p. injection and were injected with a 50 µl suspension containing $1x10^6$ primary human hepatocytes via the spleen. Eight weeks after the surgery, the humanized mice with serum human albumin levels above 0.5 mg/ml were used in experiments. For dietary treatments, the humanized mice were allowed free access to food (AL, harvested around 9–10 am, n=4), subjected to a 24-hour food withdrawal (Fast, from ~9:00 am to 9:00 am, n=5), or subjected to a 24-hour food withdrawal followed by a 4-hour refeeding (Refed from ~9:00 am to 1:00 pm, n=5). The livers were collected and stored in liquid nitrogen immediately upon sacrifice. For transcription factor agonist treatments, the humanized mice were injected with DMSO (n=4), fenofibrate (n=4, 50mg/kg, Sigma-Aldrich, Cat. F6020), rosiglitazone (n=4, 10mg/kg, Sigma-Aldrich, Cat. R2408), and GW4064 (n=4, 30mg/kg, Sigma-Aldrich, Cat. G5172) by i.p. injection (2ul/g body, starting at around 9:00 am). The livers were collected after 6 hours of fasting and stored in liquid nitrogen immediately upon sacrifice.

### Identification of functionally conserved lncRNA metabolic regulators (LMRs)

*Establishing liver lncRNA annotation in humanized mice using direct RNA sequencing (DRS)*

The flow for human and mouse liver annotation identification of humanized mice DRS has been previously described[1]. Briefly, the 500 ng of fresh polyadenylated RNAs from humanized mouse livers were used, following the Oxford Nanopore Technologies (ONT) SQK-RNA002 kit protocol, which included the optional reverse transcription step recommended by ONT. The DRS libraries were loaded onto ONT R9.4 flow cells and sequenced on the GridION or MinION platform, using the standard MinKNOW/19.12.5 protocol script. The DRS data was base-called using ONT Guppy workflow (version 2.1.0) using default setting specific parameters. The resulting FASTQ files were aligned to human and mouse genome references using minimap2/2.17 with parameters "-ax splice -uf -k14". The FLAIR pipeline with some modifications was used to identify the isoform from the aligned reads (https://github.com/BrooksLabUCSC/flair). The pipeline included removal of reads with deletion length greater than 100 nt, consideration of only reads with 5' ends that overlapped with chromatin open regions, correction of splice-site boundaries using short reads splice junctions, and inclusion of only valid junctions that were supported by at least three uniquely mapping short reads[2]. Finally, FLAIR *collapse* was utilized to generate DRS isoform annotations for human and mouse, including splicing sites and sequence information.

*Identification of syntenic LncRNA pairs*

The reference annotations were downloaded from the GENCODE database (https://www.gencodegenes.org/, GENCODE v33 for human and GENCODE vM24 for mouse). GffCompare (https://github.com/gpertea/gffcompare) was used with default settings to re-annotate the DRS annotations using the public reference annotation to obtain information about non-coding regions. For the re-annotated DRS annotations, lncRNAs were filtered by retaining only RNAs that were located in non-coding RNA regions and were longer than 200 nucleotides. Homologous gene information was used to locate syntenic regions, and bedtools/2.29.0 *intersect* was used to discover the syntenic lncRNA pairs. Syntenic lncRNA pairs were identified as those located between the same neighboring orthologous genes in both humans and mice. The orthologous gene information for humans and mice was obtained from the MGI database (https://www.informatics.jax.org/homology.shtml).

*Identification of consistently regulated lncRNA pairs*

The human and mouse liver DRS annotations were used to re-quantify short-read RNA-Seq data from humanized mice using featureCounts (subread/2.0). Differential gene expression analysis was then conducted for both human and mouse genes under various metabolic treatments using DESeq2/3.1.0 (Fast vs. AL, Fast; Refed vs. Fast, Refed; FXR agonist vs. DMSO, FXR; PPARα agonist vs. DMSO, PPARα; PPARγ agonist vs. DMSO, PPARγ). The detailed flow of differential analysis in short-read RNA-Seq was described as below. To identify significantly changed human and mouse syntenic lncRNA pairs in response to different treatments and include as many metabolically responsive lncRNA pairs as possible, the short-read RNA-Seq results were compared using a cutoff of |log2(fold change)| > 0.5 and p-value < 0.05 for differential gene expression in both humans and mice. These lncRNAs that showed significant changes in expression in both human and mouse samples were considered as metabolic lncRNA regulators. The syntenic lncRNA pairs were selected if they showed consistent regulation under at least one condition (Fast, Refed, FXR, PPARα, and PPARγ). For example, human and mouse syntenic lncRNA pairs that were both upregulated or both downregulated under the Fast treatment were defined as having consistent regulation.

*Identification of lncRNA pairs with similar functions*

The human and mouse lncRNAs that were part of consistently regulated syntenic pairs were subjected to Pearson correlation analysis with orthologous coding genes in humanized mice. The top 600 correlated orthologous genes for human or mouse lncRNAs, based on the significance of p-values, were then sent for Gene Ontology Biological Process (GO BP) pathway enrichment analysis using the R package clusterProfiler/3.18.0 with default settings (https://github.com/YuLab-SMU/clusterProfiler). Pathways significantly enriched for human and mouse lncRNAs were filtered by an adjusted p-value cutoff of less than 0.01. The similarities between pathways for human and mouse lncRNA pairs were evaluated using Fisher's exact test with the R package GeneOverlap/1.34.0. Pairs with an adjusted p-value less than 0.01 were considered to have similar functions. For one human to multiple mouse lncRNA pairs, only the pair with the most significant adjusted p-value was considered. The human and mouse lncRNA pairs with similar biological functions were defined as functionally conserved lncRNA metabolic regulators (LMRs).

**Adenovirus production and in vivo adenovirus administration**

The LMRs-pDONR221and Pabpc1-pDONR221 plasmids were first verified by sequencing, then used to construct pAdv5 adenovirus vectors using the Gateway LR reaction system (Thermo Fisher Scientific, Cat. 11791020). The shRNAs for mLMRs and mouse Pabpc1 were designed as listed in Supplementary Table 5. The hairpin template oligonucleotides, designed for shRNAs against specific targets, were synthesized by Integrated DNA Technologies. The oligonucleotides were then cloned into the pAD/Block-it system (Thermo Fisher Scientific, Cat. V49220) adenovirus vector according to the manufacturer’s protocols.

To produce the adenovirus, 5 μg of adenoviral plasmids were digested with PacI enzyme according to the manufacturer's protocol and then purified using the MinElute Reaction Cleanup Kit (Qiagen, Cat. 28206). The purified plasmids were transfected into HEK293A cells in a 6-well plate. After 48 hours, the cells were transferred to 10 cm dishes, and after approximately 5 days, the crude virus was collected. The crude viruses were amplified in HEK293A cells and purified by CsCl gradient centrifugation. The purified viruses were desalted using PD10 columns (Cytiva, Cat. 17085101) and titered with the Adeno-X Rapid Titer Kit (Takara, Cat. 632250). The adenoviruses were intravenously delivered to mice at a dose of $5 \times 10^8$ PFU/mouse for overexpression experiments or $1 \times 10^9$ PFU/mouse for knockdown experiments. For the LMRs rescue and Pabpc1 double overexpression assay, the same virus combinations were used as for the overexpression and knockdown experiments. To treat mice with a high fat diet (HFD), mice were fed with HFD (Research Diets Inc, Cat. D12492) for 8 weeks prior to receiving the virus injection. The mice were then kept on the HFD treatment for one more week before being sacrificed. Tissue samples for harvested after a 5-hour fasting (~9:00 am to 2:00 pm).

**Evaluation of LMRs rescue RNA-Seq**

*Evaluation of rescued pathways*

The RNA-Seq data was preprocessed as described, and gene expression levels were normalized using DESeq2 /3.1.0. Gene Set Enrichment Analysis (GSEA) was performed on the DESeq2-normalized RNA-Seq data to identify differentially expressed gene sets in the following groups: (A) mouse LMRs knockdown (mLMRs KD+pAdV5) versus control group (Lac+pAdv5) and (B) human LMRs rescue (hLMRs-OE+Lac) versus mouse LMRs knockdown control (mLMRs KD+pAdV5). The GSEA analysis of Gene Ontology (GO) term Biological Process (BP) was conducted using software provided by the Broad Institute (http://www.gsea-msigdb.org/gsea/index.jsp). Significantly enriched pathways were identified by |NES|>1.5, p-value <0.01, and adjusted p-value (FDR) <0.1. Rescued pathways were defined as those exhibiting opposite NES values in the mouse LMRs knockdown group (A) and the human LMRs rescue group (B), indicating rescue of pathway signatures by hLMRs overexpression. The percentage of rescue was calculated as the proportion of pathways rescued by hLMRs over the total number of significantly altered pathways in the mLMRs knockdown group.

*Evaluation of rescued DEGs*

Differential expression analysis of DEGs was performed using DESeq2/3.1.0 with default parameters. Genes with log2 (fold change) > 0.5 and p-value < 0.05 were identified as differentially expressed. To evaluate the rescue status, the commonly changed genes in mLMRs knockdown and hLMRs rescue groups were compared, with those showing opposite regulation patterns defined as rescued genes. The relative expression levels (z-score normalized) of the identified genes were then visualized using the heatmap function from the circlize/0.4.15 package in R. The enriched GO term pathways of the rescued genes were analyzed using the R package clusterProfiler/3.18.0 with the default settings.

**RNA extraction, qPCR and short-read RNA-Seq analysis**

To extract RNAs, the Trizol method was used followed by purification using the MagMAX RNA extraction kit (Thermo Fisher Scientific, Cat.AM1830). Illumina TruSeq RNA sample Prep kit was used for construction of strand-specific sequencing libraries, and sequencing was performed at the NHLBI DNA Sequencing and Genomics Core. For reverse transcription, SuperScript III First-Strand Synthesis system (Invitrogen, Cat. 11752250) was used with 500ng of RNA. Quantitative real-time RT-PCR was performed using a ViiA 7 Real-Time PCR System (Applied Biosystems Inc.) with Actb as the internal control for detecting the expressions of mouse genes in mice (C57BL/6). The PCR program was 2 minutes 30 seconds at 95°C for enzyme activation, followed by 40 cycles of 15 seconds at 95°C, and 1 minute at 60°C. Melting curve analysis was performed to confirm the real-time PCR products. The full primer sequences used are provided in Supplementary Table 5.

To analyze the humanized mice short-read RNA-Seq data, the same method as described previously was used[3]. The original FASTQ read files were first trimmed and cleaned using fastp/0.23.2 and assessed for quality using FastQC/0.11.8. A custom genome reference index was created by combining human (GRCh38.p13) and mouse (GRCm38.p6) genome sequences from the GENCODE database, and the primary assemblies were obtained from the same source. The trimmed reads were then aligned using the default settings of HISAT2/2.2.1.0 and quantified using featureCounts (subread/2.0) with public reference annotation (GENCODE v33 for human, GENCODE vM24 for mouse), as well as isoform annotation generated from the nanopore direct RNA-seq (DRS). The raw count files for both human and mouse were then imported into the R package DESeq2/3.1.0 for differential expression gene (DEG) analysis, using a log2 (fold change) cutoff greater than 0.5 and a p-value less than 0.05 to identify differential expression in all liver samples. For the analysis of mouse short-read RNA-Seq data, the FASTQ read files were first trimmed and cleaned using fastp/0.23.2 and assessed for quality using FastQC/0.11.8. The reads were then aligned using HISAT2/2.2.1.0 with an index created using the GRCm38.p6 genome. RNA-Seq aligned reads sorting, quantification (to GENCODE vm24), and DEG analysis were conducted in the same way as the humanized mouse RNA-Seq analysis. A log2 (fold change) cutoff greater than 0.5 and a p-value lower than 0.05 were used to identify DEGs.

**Analysis of public datasets**

The raw RNA-Seq data of 137 human liver samples were downloaded from the GTEx database (dbGaP, phs000424.v8.p2). The raw diversity outbred mouse liver samples treated with standard chow (n=98) or high fatty diet (n=94) were downloaded from BioProject (PRJNA295057). The raw RNA-Seq data of 14 healthy and 15 nonalcoholic fatty liver (NAFLD) patients were downloaded from PRJNA523510. The raw RNA-Seq data were analyzed as previously described, with the trimmed and cleaned reads aligned to either the human or mouse genome and quantified to reference annotations (GENCODEv33 for human, GENCODEvM24 for mouse) or nanopore DRS annotations.

The processed RNA-Seq data and protein expression data of human liver cancer were downloaded from The Cancer Genome Atlas (https://portal.gdc.cancer.gov/). The liver samples were stratified into high and low expression groups based on the RNA expression levels of hLMR1 or PABPC1, with the top quartile representing the high expression group and the bottom quartile representing the low expression group. The relative total protein expression levels in each sample were calculated as the sum of all protein expression levels.

**Human- and mouse-LMRs clone, gel analysis and verification**

To clone human- and mouse-LMRs, the Gateway BP cloning system (Thermo Fisher Scientific) was used in accordance with the manufacturer's instructions. The attB PCR primers were designed by adding the attB sequence to the 5'-end of the forward (GGGGACAAGTTTGTACAAAAAAGCAGGCT) and reverse (GGGGACCACTTTGTACAAGAAAGCTGGGT) primers (Supplementary Table 5). The human- and mouse-LMRs DNA fragments were amplified by PCR from humanized liver cDNA samples. Following gel electrophoresis, the attB-flanked PCR products with the appropriate sizes were purified and then cloned into pDONR221 vectors using the Gateway BP reaction system. Sequencing by Synthesis (SBS) sequencing was employed for the identification of the novel gene sequences, which were then compared to the nanopore DRS annotation using BLAST (https://blast.ncbi.nlm.nih.gov/Blast.cgi).

**In vitro translation**

In vitro translation was carried out using the TnT Quick Coupled Transcription/Translation System from Promega (Cat. L1170), as per the manufacturer's instructions. The translated proteins were detected by IRDye streptavidin (LI-COR), which binds biotinylated lysine incorporated into the translated proteins. An open reading frame of luciferase DNA provide by manufacturer was used as a positive control.

**RNA pulldown assay**

As previously described, an RNA pulldown assay was performed[3, 4]. In brief, biotin-labeled RNA was produced using T7 RNA polymerase (Invitrogen, Cat. AM2085) and biotin RNA labeling mix (Roche, Cat. 11685597910) and purified using the RNeasy Mini Kit (QIAGEN, Cat. 74004). As a control, the hLMR1 reverse direction sequence (hLMR1 anti-sense) was employed. Five mg of precleared humanized mouse liver lysate (supplemented with 0.2 mg/mL heparin, 0.2 mg/mL yeast tRNA, and 1 mM DTT) was combined with 3 μg of folded RNA before being incubated at 4°C for an hour. After that, each binding reaction received 60 μL of washed streptavidin-coupled Dynabeads (Invitrogen, Cat 65306), which were then incubated for an additional hour at 4°C. The beads were washed five times with lysis buffer (1 mM EDTA, 20 mM Tris pH 7.4, 150 mM NaCl, 0.5% Triton X-100 with Protease/Phosphatase Inhibitor Cocktail (Thermo Fisher Scientific, Cat. 78429) and RNaseOUT (Thermo Fisher Scientific, Cat. 10777019) and then heated in 1× lithium dodecyl sulfate (LDS) loading buffer at 70°C for 10 minutes. The retrieved proteins were visualized by SDS-PAGE and silver staining (Invitrogen, Cat. LC6070), and the unique protein bands shown in the hLMR1 RNA pulldown were identified by mass spectrometry analysis at NHLBI Proteomics Facility. Western blotting assay was used to verify the binding of PBPC1

(Invitrogen, Cat. PA5-17599) with hLMR1 and mLMR1. hLMR1 and mLMR1 antisense as well as a beads-only sample were used as negative controls.

**RNA immunoprecipitation (RIP)**

RIP analysis was performed as described previously [3]. Frozen humanized liver tissues were homogenized in RIP buffer (pH 7.4, 1 mM EDTA, 20 mM Tris, 150 mM NaCl, 0.5% Triton X-100 with Protease/Phosphatase Inhibitor Cocktail and RNaseOUT) using a Dounce homogenizer for 15–20 strokes. For each RIP, 30 μL of washed Dynabeads coupled with Protein G were incubated with 5 μg of rabbit IgG or anti-PABPC1 antibody (Invitrogen, Cat. PA5-17599) in a 300 μL solution of RIP buffer containing 0.2 mg/mL heparin, 0.2 mg/mL BSA and 0.2 mg/mL EcoRI tRNA for one hour. The antibody-coupled beads were then added to a 500 μL solution of RIP buffer containing 5 mg of liver tissue lysates and incubated for 3 hours at 4°C with gentle rotation. The beads were washed briefly five times with RIP buffer; during the final wash, one-fifth of the beads were used for protein analysis, while the remaining beads were resuspended in 1 mL of TRIzol for RNA extraction. The coprecipitated RNAs were isolated and subsequently analyzed by RT-PCR.

**RNA dot-blot assay**

Biotinylated full-length RNA of hLMR1 and mLMR1 (0.5-1 μg) were incubated with recombinant c-myc-tagged PABPC1 proteins (50 ng, OriGene Technologies, Cat. TP307354) in the RNA-protein binding buffer (50 mM Tris-HCl pH 7.9, 10 mM β-ME, 10% glycerol, 5 mM MgCl2, 100 mM KCl, 0.1% NP-40, 10 mg/mL yeast tRNA, 40 U/μL RNase inhibitor) on ice for one hour. Then, the binding mixture was transferred into a pre-cooled 96-well round-bottomed plate and placed on ice with an ultraviolet (UV) crosslinker (Analytik Jena, CL-3000), and 150 mJ/cm2 radiation was applied for 10 minutes. After UV-crosslinking, the RNAs were partially digested by RNase A (Thermo Fisher, Cat. AM2271) at a 1:50 ratio for 30 minutes at 37°C. Then, RNA-protein complexes were subjected to purification by c-myc tag magnetic beads (Thermo Fisher, Cat. 88842), and purified RNA-protein complexes were treated with proteinase K (Thermo Fisher, Cat. 25530049), which removed proteins but left the protected RNAs. The RNA samples were dissolved in ice-cold 10mM NaOH and 1mM EDTA buffer, and the Nitrocellulose membranes (LI-COR, Cat. 46639480) were baked at 80°C for 2 hours before use. Then, 5-10 μL of biotin-

RNA samples were dropped onto the membrane for each well with a Bio-Dot apparatus (BIO-RAD, Cat. 706545), avoiding touching the membrane with the pipette tip, and incubated with DNA probes (30-mer antisense DNA oligonucleotides, incubation buffer: 6X SSC buffer with 0.5% SDS and 1mM EDTA) at 65°C for 2 hours. Then, the hybridized membrane was washed with washing buffer (2X SSC buffer with 0.5% SDS) at 37°C, 50°C, and 65°C for 20 minutes each. Finally, the protein-bound RNA sequence was visualized by detecting Streptavidin-AlexaFluor® 800 (IRDye-800CW Streptavidin, LI-COR, Cat. C41209-03) signals. The DNA probes, which were synthesized at Integrated DNA Technologies (IDT), are listed in Supplementary Table 5. The schematic diagram of the assays can be found in Supplementary Figure 6*D*.

**Competition RNA pulldown assay**

For the full-length RNA competition assay of hLMR1 and mLMR1, equimolar amounts of biotin-labeled mLMR1 and non-biotin labeled hLMR1 (or biotin-labeled hLMR1 and non-biotin labeled mLMR1) were incubated with precleared humanized mouse liver lysate, with the same procedure used for the RNA pulldown assay. For the hLMR1-dot24 RNA (CCAACAUGGUGAAACCCUGUCUCUACUAAA) competition with full length mLMR1 or the mLMR1-dot30 RNA (CCAGGGCCAAGAAGUGGGAGUGGGUGGGUA) competition with full length hLMR1, we prepared four concentration gradients (biotin-labeled mLMR1:hLMR1-dot24 RNA oligo=1:1, 1:5, 1:10, 1:15 or biotin-labeled hLMR1:mLMR1-dot30 RNA oligo=1:1, 1:5, 1:10, 1:15) based on the number of moles and incubated them with precleared humanized mouse liver lysate. The subsequent steps were the same as described for the RNA pulldown assay. The samples were then subjected to electrophoresis on 4–12% gradient PAGE gels (Thermo Fisher, Cat. NP0335BOX) and immunoblotting with anti-PABPC1 antibody to detect the signals. The RNA oligos were synthesized by IDT company. The schematic diagram of the assays can be found in Supplementary Figure 6*C* and *E*.

**Immunoblotting**

For Immunoblotting analyses, liver tissues were lysed in 1% SDS lysis buffer containing phosphatase inhibitors (Sigma) and a protease inhibitor cocktail (Roche). The lysate was subjected to SDS–PAGE, transferred to polyvinylidene fluoride (PVDF) membranes, and incubated with the primary antibody followed by the fluorescence conjugated secondary antibody (LI-COR). The bound antibody was visualized using a quantitative fluorescence imaging system (LI-COR). The following antibodies were used: Scd1 antibody (Cell Signaling Technology, Cat. 2794, Dilution: 1:1000), Fasn (Cell Signaling Technology, Cat. 3180, Dilution: 1:1000), Acaca (Cell Signaling Technology, Cat. 3676, Dilution: 1:1000), mTOR antibody (Cell Signaling Technology, Cat. 2983, Dilution: 1:1000), Phospho-mTOR (Ser2448) Antibody (Cell Signaling Technology, Cat. 2971, Dilution: 1:1000) and Alpha-Tubulin antibody (Sigma-Aldrich, Cat. MABT522, Dilution: 1:1000).

**Design of PABPC1 deletion mutants and RIP assay**

The cDNAs encoding human PABPC1 ORF (NCBI Reference Sequence: NM_002568.4) were cloned into the BglII and KpnI sites of the plasmid p-3xFLAG-CMV-13 vector. The construction of the five PABPC1 deletion mutants, including pMLLE (1-370aa), pRRM1 (99-636aa), pRRM1/2 (191-636aa), pRRM1/2/3 (294-636aa), and pRRM1/2/3/4 (542-636aa), were performed by inserting the DNA fragments with BglII and KpnI sites into the C-terminal p-3xFLAG-CMV-13 vector. All the plasmids were verified by sequencing. For the RIP assay, human primary hepatocytes (Lonza, Cat. HUM181001B) or mouse primary hepatocytes (Cell Biologics Inc., Cat. C57-6224F) were transfected with PABPC1 full length and deletion mutants FLAG-tag vectors. After 36-48 hours of transfection, cells were lysed in RIPA buffer (Boston Bioproducts, Cat. BP-115) containing protease/phosphatase inhibitor Cocktail (Thermo Fisher, Cat. 78440) and RNase inhibitors (Thermo Fisher, Cat. 10777019). The lysate was placed on ice for 15-20 minutes and then centrifuged at 4°C for 10 minutes, and the supernatant was used for immunoprecipitation using anti-Flag antibody (Sigma-Aldrich, Cat. F1804-200UG). The following steps were the same as previously described[3]. The primers for the qPCR are listed in Supplementary Table 5.

**Prediction of RNA structure**

The secondary structures of hLMR1-dot24 RNA (CCAACAUGGUGAAACCCUGUCUCUACUAAA) and mLMR1-dot30 RNA (CCAGGGCCAAGAAGUGGGAGUGGGUGGUA) were predicted by RNAfold using the default settings (http://rna.tbi.univie.ac.at/cgi-bin/RNAWebSuite/RNAfold.cgi). The models displaying the minimum free energy (MFE) structure, along with encoded base-pair probabilities, were presented. The tertiary structures of the RNA motifs were predicted by Alphafold3 using the default setting[5] and were visualized by ChimeraX/1.6.

**Chromatin immunoprecipitation (ChIP)**

The mouse livers were minced, washed twice with PBS, and incubated with 1.5% (w/v) formaldehyde for 20 minutes for crosslinking. Chromatin samples were subsequently treated with 1X glycine for 5 minutes to quench the crosslinking. Chromatin immunoprecipitation (ChIP) was conducted following the protocol outlined in the SimpleChIP Plus Enzymatic Chromatin IP Kit (Cell Signaling, Cat. 9005). In brief, the chromatin samples were sonicated and then incubated with 2 µg of antibodies specific to SREBP1, RNA polymerase II, or control IgG overnight at 4°C. The immunoprecipitated chromatin was applied to Protein G Magnetic Beads for 1 hour. The beads were washed three times with 1X ChIP buffer and once with 1X ChIP buffer containing 350 mM NaCl. Subsequently, the beads were incubated overnight at 65°C to reverse the crosslinking. DNA was isolated and quantified using quantitative polymerase chain reaction (qPCR). SREBP1 binding peaks were identified based on a published SREBP1 ChIP-Seq data [6]. Primer sequences are provided in Supplementary Table 5.

**Oil Red O Staining**

The staining of lipid droplets in liver tissues were performed according to the manufacturer's protocols (VitroVivo Biotech, Cat. VB-3007). Frozen liver sections were cut at 8 to 10 mm and air-dried onto slides. The sections were fixed in 10% buffered formalin for 20 minutes at room temperature and briefly washed with running tap water for 3 minutes. Next, the sections were placed in Pre-Stain Solution for 2 minutes followed by staining in pre-warmed Oil Red O solution for 10 minutes at room temperature. After staining, the sections were differentiated in Differentiation Solution for 5 minutes at room temperature and then rinsed in 2 changes of distilled water at room temperature. The nuclei were lightly stained with Mayer's Hematoxylin Solution for 1 minute and the sections were rinsed with tap water for 3 minutes. Finally, the sections were mounted with aqueous mounting medium. The stained slides were immediately imaged under a microscope and the lipid droplets were quantified using ImageJ. The proportions of lipid droplet area to the total image area were calculated and standardized to the control group.

**Measurement of plasma and liver triglyceride (TG)**

The TG levels were measured according to the manufacturer's directions. For the detection of plasma TG, fresh blood was immediately centrifuged in a refrigerated centrifuge (4°C, 12000 rpm/min, 10 min), and the supernatants were transferred to new Eppendorf (EP) tubes for immediate testing. The remaining plasma samples were stored at -80°C. For the detection of liver TG, approximately 30mg of frozen tissue powder was weighed and added to 2ml round-bottomed EP tubes. The 20 µl of 5% NP 40 was added per mg of tissue and homogenized using the TissueLyser LT (Qiagen, Cat. 85600). The samples were slowly heated to 90°C in a water bath and incubated for 2-5 minutes, then slowly cooled to room temperature. This step was repeated once, and the samples were centrifuged at 12000 rpm/min for 10 min. The supernatants were transferred to new EP tubes and mixed thoroughly using pipettes, then immediately tested. For TG detection, 5 µl of plasma or 8 µl of liver TG supernatants were mixed with an equal volume of glycerol standard solution (Sigma-Aldrich, Cat. G7793) and blank controls (PBS for plasma, 5% NP40 for liver) and added to 80 µl of free glycerol reagent (Sigma-Aldrich, Cat. F6428-40ML). The mixtures were shaken well and incubated at 37°C for 5 minutes, then the absorbances were measured at 540 nm. 20 µl of triglyceride reagent (T2449-10ML) was added to each well, shaken thoroughly, incubated at 37°C for 5 minutes, and the absorbances were measured at 540 nm. TG relative levels were calculated according to the manufacturer's instructions.

**Polysome fractionation analysis**

The polysome fractionation analysis was conducted following a previously described method[7]. Stock solutions of 10%, 20%, 30%, 40%, or 50% sucrose were prepared and stored at 4°C. The sucrose gradients were prepared a day in advance of the experiment, and the SW41Ti rotor and buckets were pre-cooled at 4°C overnight. Using the prepared sucrose solutions, 2.2 ml of 10% sucrose gradient was added to the bottom of a thin-wall tube, and subsequent layers (2.2 ml each of 20%, 30%, 40%, and 50%) were underlaid using a Pasteur pipet and manual pipettor. The gradients were left to equilibrate at 4°C overnight to ensure a linear gradient. Approximately 100 mg of frozen liver powder was weighed and added to 1.5 ml of lysis buffer containing 100 mg/ml cycloheximide (CHX, Sigma-Aldrich, Cat. C7698) and RNAseOut. The samples were gently homogenized 20-25 times using a Dounce homogenizer on ice and then transferred to new nuclease-free 1.5 ml Eppendorf (EP) tubes. After incubation on ice for 10 minutes with occasional tube inversion every two minutes, the samples were centrifuged at 12,000 rpm/min in a refrigerated centrifuge for 10 minutes, and the supernatants were carefully transferred to new 1.5 ml EP tubes (650 µl each). The supernatants were then layered onto the top of 10-50% sucrose gradients, and the gradients were centrifuged for 90 minutes in a SW41Ti swinging bucket rotor at 190,000 x g (~39,000 rpm) at 4°C with maximum acceleration and brake using a Beckman ultracentrifuge. The RNA content of each fraction was measured using a 254 nm spectrophotometer at a speed of 1 ml/min fractions with a fractionating system (Brandel, Cat.BR-188). Polysome fractions in the gradients were defined based on the distribution of RNA as previously described [7].

**Free amino acid analysis**

The snap-frozen liver was finely powdered in liquid nitrogen. Samples from the same groups were equally pooled, and 20 mg of frozen liver powder was weighed and dissolved in 400 µl of cold 100% methanol (MeOH) in a new round-bottom Eppendorf (EP) tube. Subsequently, 400 µl of double-distilled water was added, along with a stainless-steel bead (Qiagen, Cat. 69989). Homogenization was performed using the TissueLyser (Qiagen, Cat. 85600) for 4 minutes at 50 Hz. The tubes were then placed on ice for 15 minutes, and 400 µl of cold chloroform and norvaline (to a final concentration of 1 nmol/µl) were added. The samples were shaken at 1000 rpm/min for 20 minutes at room temperature. Vigorously vertexing for 2 minutes was followed by centrifugation at 16,000 g for 10 minutes at 4°C. Subsequently, 300 µl of the upper phase was transferred to a new 1.7 ml EP tube. The extracted material was dried using a SpeedVac. The resulting pellet was dissolved in 50 µl of water. Centrifugation at 16,000 g for 10 minutes at 4°C was then performed, and the supernatant was transferred to a sample vial (Agilent, Cat. 5188-6591) for High-Performance Liquid Chromatography (HPLC) on an Agilent 1100 system with an amino acid analysis column (Agilent, Cat. 959963-302). The HPLC procedure followed the manufacturer's instructions and was analyzed using ChemStation/B.04.03. The relative expression levels of each amino acid were normalized to the internal control norvaline expression levels.

**Quantification and statistical analysis**

The figure legends provide information on the number of mice utilized for each experiment. Every experiment was repeated a minimum of three times or involved a minimum of three distinct biological replicates. Data are presented as mean ± standard error of the mean (SEM). Unless specified otherwise, statistical significance in mean values between two populations was assessed using a two-tailed Student's t-test (*$P < 0.05$; **$P < 0.01$; ***$P < 0.001$; # $P < 0.05$; ## $P < 0.01$; ###$P < 0.001$). GraphPad Prism 9 software and R/4.3.1 were employed for statistical analysis and figure generation.

### Study approval

All animal experiments were performed in accordance with and with approval from the NHLBI Animal Care and Use Committee or the Animal Care Committee of the CIEA, Kawasaki, Japan. All human-related data sets were downloaded from public databases.

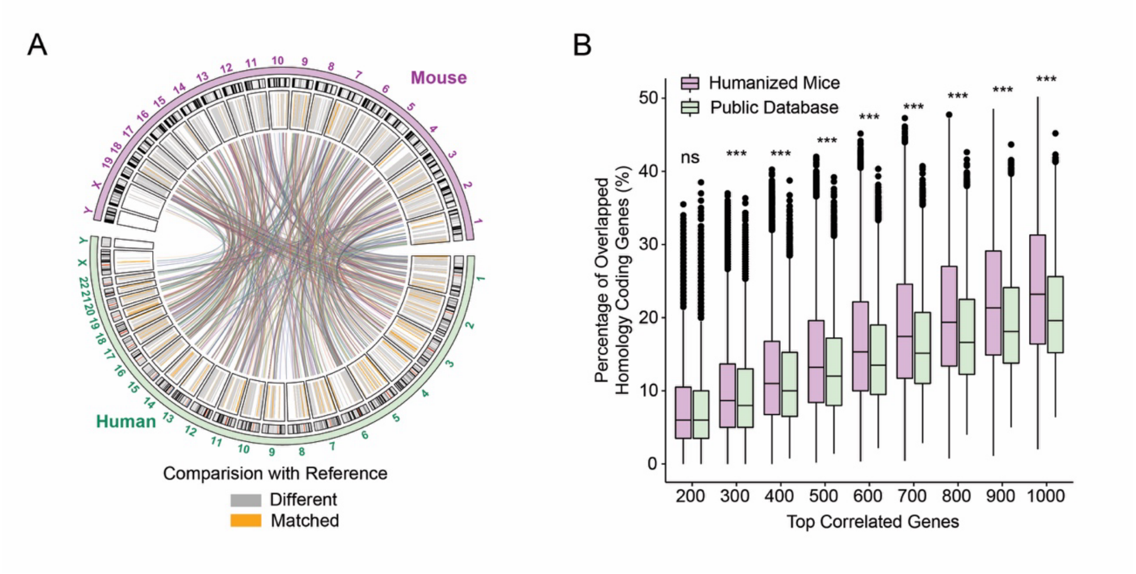


**Supplementary Figure 1 A function-based strategy to identify functionally conserved lncRNA metabolic regulators (fcLMRs) between humans and mice**

(A) Circular diagram of the genomic locations of human and mouse lncRNAs. The isoform of the lncRNA differing from the reference genome was marked in grey on the inner circle, while orange indicated that they were the same. LncRNA pairs located in the same syntenic regions in humans and mice were represented by the central line. (B) Evaluation of the efficacy to identify homologous coding genes between human and mouse by correlation analyses using gene expression data of humanized mice or public database. We first downloaded the list of all homologous protein-coding genes in humans and mice from MGI database (https://www.informatics.jax.org/homology.shtml) and performed Pearson correlation to calculate the correlation coefficients and p-values of each human or mouse gene with all remaining genes within the same species. The correlation analyses were performed with human and mouse RNA-seq data from humanized mice or public datasets. The humanized liver RNA-Seq included all treatments (n=34). Public datasets included the human liver RNA-seq dataset from GTEx database (n=137) and the outbred mouse liver RNA-seq dataset PRJNA295057 (standard chow, n=98). Based on the significance of the p-values, all genes correlated with each gene were ranked and the number of top genes were selected based on the listed cutoff (200-1000). We then calculated how many selected top genes for a homologous gene pair are overlapped and expressed them as a percentage for each cutoff. We then presented the average percentage of all gene pairs as bar plots in humanized mice and public databases. The higher average percentage indicated the analysis is more effective in identifying known homologous protein-coding gene pairs. ***p<0.001, two-tailed unpaired Student's t-test.

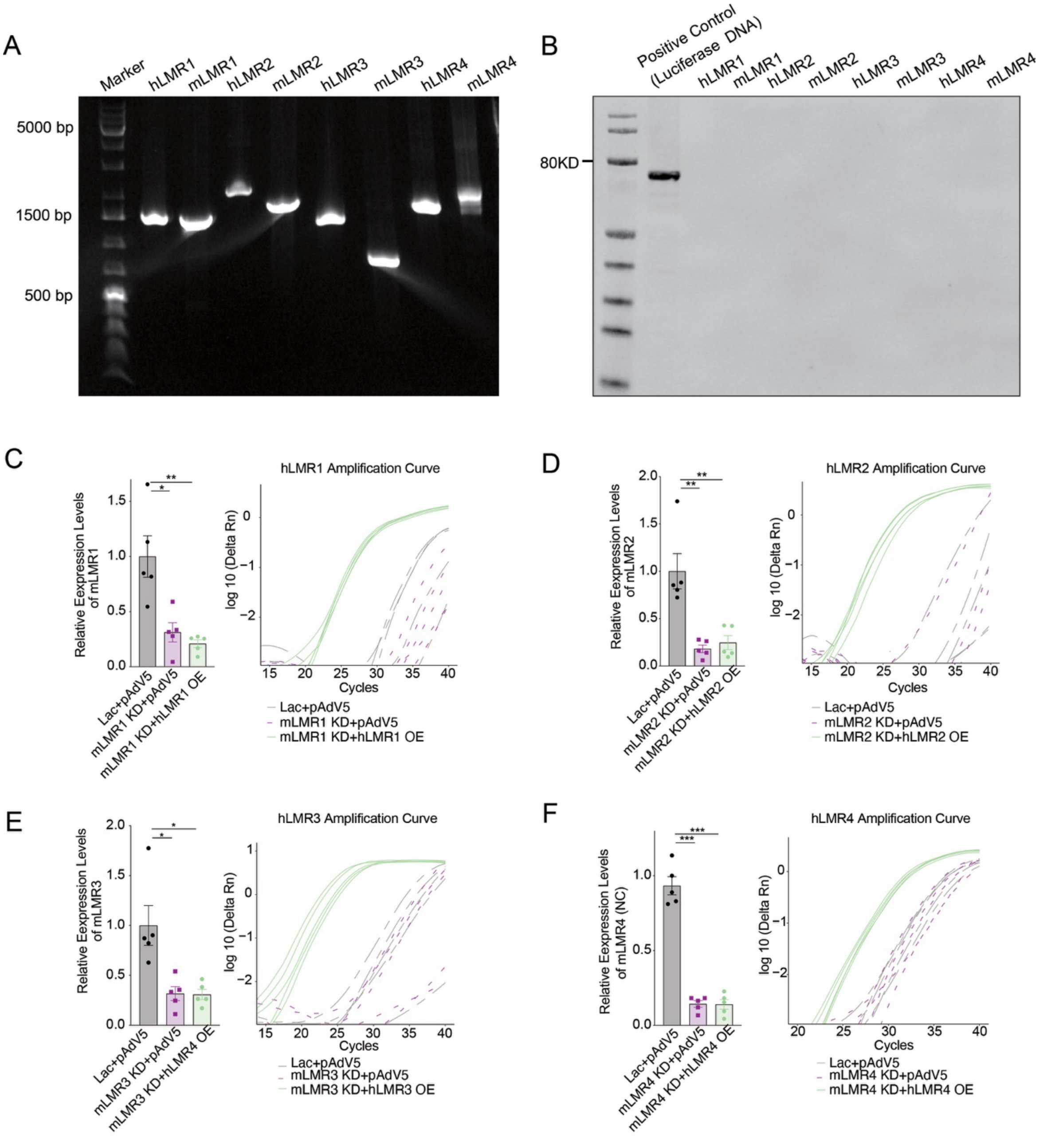


**Supplementary Figure 2 Characterization of hLMRs and mLMRs**

(A) Gel analysis of the PCR cloned hFCL1-4 and mFCL1-4. (B) In vitro translation analyses of hFCL1-4 and mFCL1-4. The luciferase protein DNA was used as positive control. (C-F) Mice were injected with three groups of adenoviruses (n=5 per group), including control viruses for knockdown and overexpression (Lac+pAdv5, control group), mLMR knockdown viruses and control viruses for overexpression (mouse-lncRNA KD+pAdv5, mLMR knockdown group), and mLMR knockdown viruses and hLMR overexpression group (mLMR KD+hLMR OE, hLMR rescue group). The expression levels of mLMR1-4 and amplification curves of hLMR1-4 were analyzed by qPCR and normalized to control group. As hLMR1-4 were not expressed in mouse livers, their amplification curves instead of relative expression levels were shown.

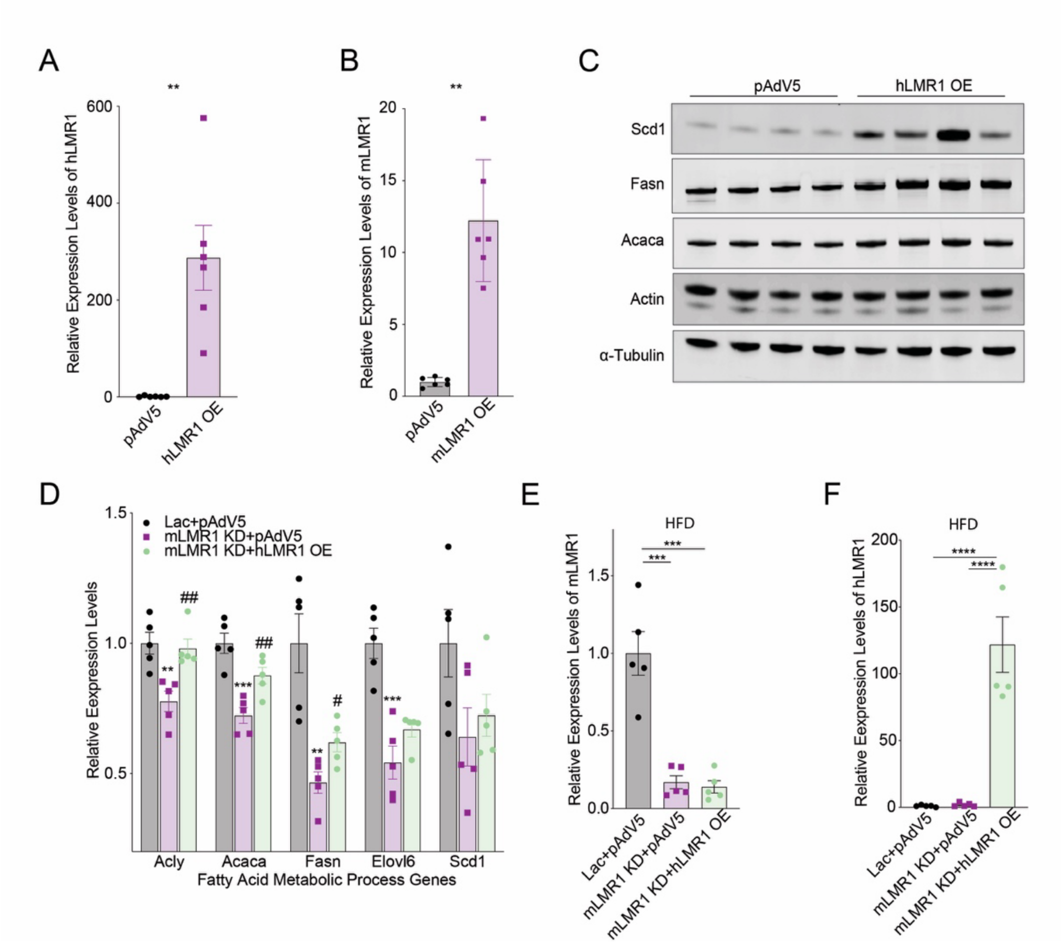


**Supplementary Figure 3 hLMR1 and mLMR1 modulate lipid metabolism**

(A and B) The relative expression levels of hLMR1 (A) or mLMR1 (B) in their overexpression mice. (C) Immunoblotting analysis of the expression levels of Scd, Fasn, Acaca, Actin, and α-Tubulin in mice subjected to hLMR1 overexpression (hLMR1 OE) and pAdV5 (n=4 per group). (D) The relative expression levels of representative genes in the fatty acid metabolic process in mice receiving the following adenovirus combinations: Lac+pAdv5, mLMR1 KD+pAdv5, and mLMR1 KD+hLMR1 OE. Data represent mean ± SEM. *p<0.05, **p<0.01, ***p<0.001, represent a comparison between mLMR1 KD+pAdv5 and Lac+pAdv5, two-tailed unpaired Student's t-test. $^{\#}$p<0.05, $^{\#\#}$p<0.01 represent a comparison between mLMR1 KD+hLMR1 OE and mLMR1 KD+pAdv5, two-tailed unpaired Student's t-test. (E and F) The relative expression levels of mLMR1 (E) and hLMR1 (F) in HFD-fed mice receiving the following adenovirus combinations: Lac+pAdv5, mLMR1 KD+pAdv5, and mLMR1 KD+hLMR1 OE.

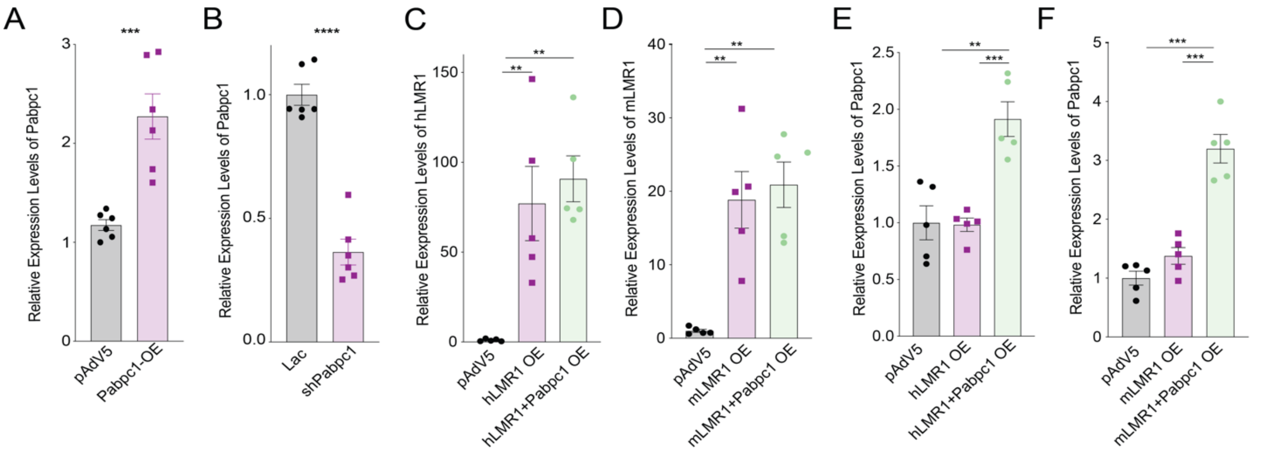


**Supplementary Figure 4 hLMR1 and mLMR1 modulate lipid metabolism and translation via PABPC1**

(A and B) The relative expression levels of Pabpc1 in mice with Pabpc1 overexpression (A) (n=6 per group) and knockdown (B) (n=6 per group). Data represent mean ± SEM, ***p<0.001, ****p<0.0001, two-tailed unpaired Student's t-test. (C-F) The relative expression levels of hLMR1 (C), mLMR1 (D), and Pabpc1 (E-F) in mice (n=5 per group) receiving the following adenovirus combinations: pAdV5 (control group), human/mouse LMR1 overexpression (h/mLMR1 OE), and co-expression of human/mouse LMR1 and Pabpc1 (h/mLMR1+Pabpc1 OE) adenovirus. Data represent mean ± SEM, **p<0.01, ***p<0.001, two-tailed unpaired Student's t-test.

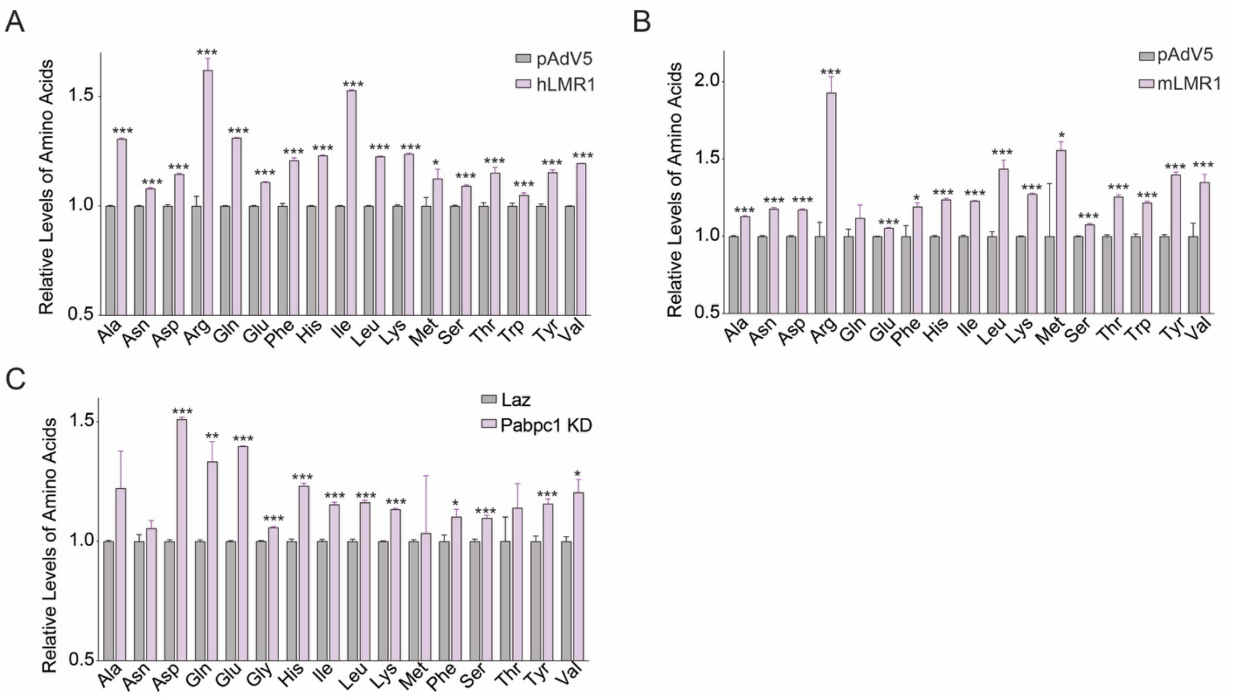


**Supplementary Figure 5 h/mLMR1 interact with Pabpc1 to modulate levels of amino acids**
(A-C) Relative levels of amino acids in the livers of mice receiving one of the following adenoviruses: hLMR1 (A), mLMR1 (B), or Papbc1knock-down (KD) (C). Liver samples from the same group were pooled, and amino acid levels were measured by HPLC in triplicate. *p<0.05, **p<0.01, ***p<0.001, two-tailed unpaired Student's t-test.

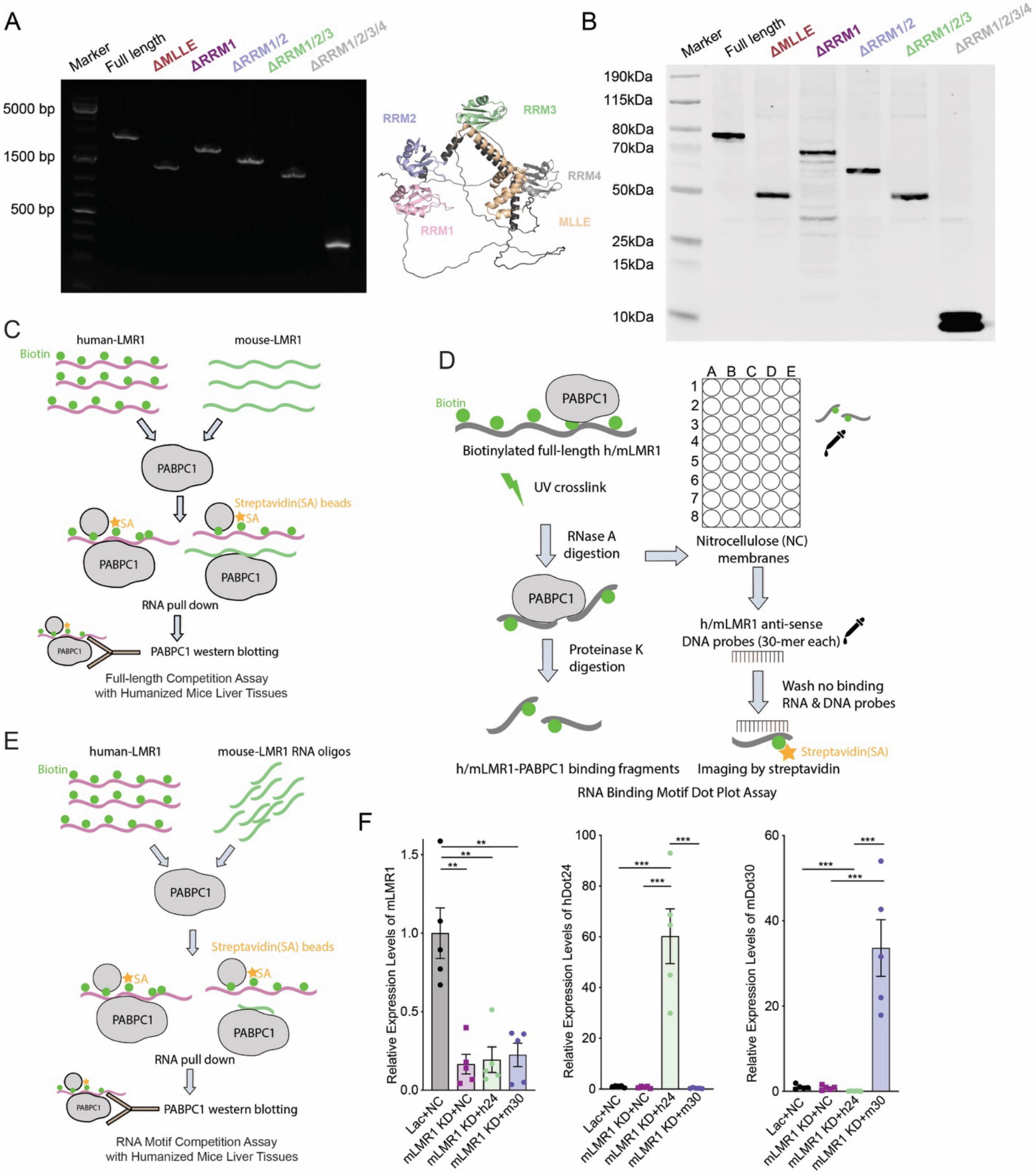


**Supplementary Figure 6 Similar functions of hLMR1 and mLMR1 are mediated by conserved PABPC1 docking**

(A) Gel analysis of the PCR cloned PABPC1 variants with deletions of functional domains (left). The 3D structure of PABPC1 with functional domains labeled (right). (B) Western blotting was performed to determine the protein expression levels of different PABPC1 domain deletion constructs in 293A cells. The cells were transfected with various PABPC1 domain deletion constructs, and after 48 hours, protein extracts were collected and the expression levels of PABPC1 with different domain deletions were detected using a flag antibody. (C-E) Schematic diagram of the LMR1 full-length competition assay (C), LMR1 RNA binding motif dot plot assay (D), LMR1 RNA binding motif competition assay (E). (F) The expression levels of mLMR1, hLMR1 fragment 24 (h24), and mLMR1 fragment 30 (m30) in mice receiving the following adenovirus combinations: Lac+NC (Negative control), mLMR1 KD+NC, mLMR1 KD+h24, and mLMR1 KD+m30, relatively. Data represent mean ± SEM, $^{**}p<0.01$, $^{***}p<0.001$, two-tailed unpaired Student's t-test.

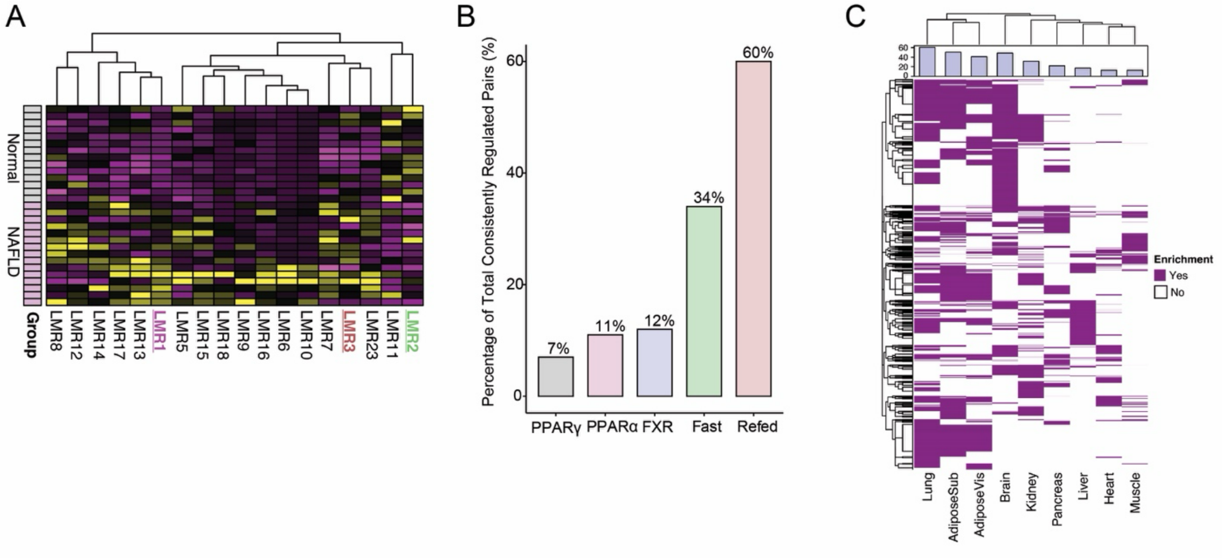


**Supplementary Figure 7 Distribution of lncRNAs in different treatment groups and tissues**

(A) Heatmap of gene expression levels of human LMRs that were significantly altered in fatty liver. The differentially expressed genes (DEGs) were defined as | log2 (fold change) | > 0.5 and p value 0.05. The gene names of interest were marked with different colors on the heatmap. LMR1 was marked in purple, LMR2 in green, and LMR3 in deep red. (B) The percentages of syntenic lncRNAs with conserved regulation distributed across different treatment conditions. (C) Heatmap of human lncRNA expression enrichment in major human tissues. The percentage of lncRNA enrichment in each tissue is summarized at the top.
